# AFC kinases regulate warm temperature-responsive growth in *Arabidopsis*

**DOI:** 10.1101/2024.06.21.600040

**Authors:** Benjamin Dimos-Röhl, Felix Ostwaldt, Jannik Bäsmann, Paula Hausmann, Philipp Kreisz, Markus Krischke, Ruth Lintermann, Christoffer Lutsch, Philipp C. Müller, Daniel Schubert, Miriam Strauch, Christoph Weiste, Tingting Zhu, Ive De Smet, Florian Heyd, Daniel Maag

**Affiliations:** Laboratory of RNA Biochemistry, Institute of Chemistry and Biochemistry, Freie Universität Berlin, 14195 Berlin, Germany; Department of Pharmaceutical Biology, Faculty of Biology, Julius-von-Sachs-Institute of Biosciences, Julius-Maximilians-Universität Würzburg, 97082 Würzburg, Germany; Epigenetics of Plants, Institute of Biology, Freie Universität Berlin, 14195 Berlin, Germany; Department of Plant Biotechnology and Bioinformatics, Ghent University, 9052 Ghent, Belgium; VIB Center for Plant Systems Biology, 9052 Ghent, Belgium

**Keywords:** alternative splicing, post-transcriptional regulation, thermomorphogenesis

## Abstract

Plants respond to elevated temperatures with enhanced elongation growth that depends on rapid transcriptional, post-transcriptional, and post-translational reprogramming. However, it is unclear how temperature information integrates with the splicing machinery to establish warm temperature-dependent splicing patterns. In animals, CDC2-LIKE KINASES (CLKs) function as body temperature sensors that control temperature-dependent splicing via phosphorylation of serine/arginine-rich (SR) proteins. Here we demonstrate that the CLK-homologous ARABIDOPSIS FUS3-COMPLEMENTING (AFC) kinases likewise regulate post-transcriptional RNA processing to control warm temperature-dependent growth in Arabidopsis. The contrasting temperature-activity profiles of the three AFCs depend on specific structural elements, including a conserved activation segment within the kinase domain. Combining protein structure prediction with site-directed mutagenesis, we provide insights into structural features that determine the different temperature-activity profiles of the three AFC paralogs. Analyses of *afc* mutant plants demonstrate their role in establishing temperature-dependent splicing patterns and thermomorphogenic hypocotyl elongation. Finally, our data indicate SR34 and SR34a as phosphorylation targets mediating temperature-dependent hypocotyl elongation downstream of AFCs. In conclusion, our study provides evidence that temperature-controlled AFC activity is evolutionarily conserved between plants and animals and implicates AFCs in the control of thermomorphogenesis.

## Introduction

Plants exhibit a substantial morphological and developmental plasticity in response to varying temperature conditions. Accordingly, a moderate temperature increase within the physiological range already has profound effects on plant architecture including enhanced hypocotyl, petiole, and root elongation growth as well as thermonastic leaf movement^1^. These warm temperature-dependent morphological alterations have been grouped under the term thermomorphogenesis^2, 3^. At the cellular level, thermomorphogenic growth responses are accompanied by a substantial rearrangement of the proteome that depends on epigenetic^4^, transcriptional^5^, post-translational^6, 7^ and hormonal regulation^8, 9^. The basic helix-loop-helix proteins PHYTOCHROME INTERACTING FACTOR 4 (PIF4) and PIF7 have been identified as the central transcriptional regulators in thermomorphogenesis signalling^10, 11, 12^. Upon exposure to elevated temperatures, PIF4 is derepressed and induces the transcription of temperature-responsive genes including that of auxin-biosynthetic enzymes, thereby positively regulating elongation growth^12^. Downstream of auxin, functional brassinosteroid signalling is required for thermomorphogenic growth responses involving the transcription factors BRASSINAZOLE-RESISTANT 1 (BZR1) and BZR2^3^. However, PIF4- and auxin-independent regulation of thermomorphogenesis has also been described^7^.

While the hormonal and transcriptional regulation underlying thermomorphogenesis is relatively well understood, the contribution of additional regulatory layers such as post-transcriptional regulation only came into view in recent years^13, 14^. For example, seedlings defective in the spliceosomal protein SNW/SKI-INTERACTING PROTEIN (SKIP), which display global defects in pre-mRNA splicing^15^, show reduced hypocotyl elongation growth in response to elevated temperature^14^. At the genome-wide level, it has been estimated that 30% to 60% of all intron-containing genes undergo alternative splicing in response to various high temperature treatments in Arabidopsis^16^. Thus, warm temperature-induced alternative splicing emerges as central regulatory mechanism that allows for fast adjustments of the proteome in response to changing temperature conditions^13^.

Splicing of precursor mRNA is catalysed by the spliceosome, a large complex in the nucleus consisting of five small nuclear ribonucleoprotein (snRNP) complexes and various auxiliary factors. Among them, evolutionarily conserved serine/arginine-rich (SR) proteins govern splice-site selection by the recognition of *cis*-regulatory elements within the pre-mRNA, thereby regulating both constitutive and alternative splicing^17^. SR proteins consist of one or two N-terminal RNA-binding domains and a C-terminal RS domain, which contains multiple arginine/serine dipeptide repeats^18^. The expression of SR proteins is regulated in a temperature-dependent manner at the transcriptional and post-transcriptional level^19, 20^. In addition, they possess several phosphorylation sites that determine their activity and subnuclear localisation^21^. Notably, in response to an increase in temperature several SR proteins are rapidly phosphorylated, suggesting warm temperature-dependent post-translational regulation of SR protein activity^22^.

In animals, these phosphorylation sites are targeted by CDC2-LIKE KINASES (CLKs). Recently, it has been shown that CLK1 and CLK4 regulate temperature-dependent alternative splicing in human cells within the physiological temperature range through phosphorylation of SR proteins^23^. Moreover, *in vitro* kinase assays using mouse CLK1 and CLK4 demonstrated that temperature sensitivity is an intrinsic property of these proteins that depends on subtle conformational changes of a conserved activation segment consisting of 25 amino acid within the active centre of the kinase domain^23^. Altogether, these observations suggest that, in animals, CLKs act as conserved temperature sensors that facilitate context-dependent gene expression through their regulatory effect on the splicing machinery.

The Arabidopsis genome contains three genes that are homologous to human *CLKs*, i.e., *ARABIDOPSIS FUS3-COMPLEMENTING 1* (*AFC1)*, *AFC2*, and *AFC3*^24^. Both, *in vitro* and *in vivo* data support their involvement in pre-mRNA processing through binding and phosphorylation of SR proteins^25, 26, 27, 28^. Overexpression of the tobacco AFC2 homolog in Arabidopsis led to reduced growth and late flowering, suggesting a role of AFC2 in plant development^27^. More recently, functional roles during low temperature acclimation and thermomorphogenesis have been proposed for AFC3 and AFC2, respectively^26, 29^. However, the mechanistic link between AFC activity, RNA processing and temperature responsiveness remained elusive. Moreover, it is unknown whether plant AFC kinases, alike their animal counterparts, also possess thermosensory properties.

In the present study, we addressed the molecular mechanisms and structural details underlying temperature-controlled AFC activity and assessed their functional role in thermomorphogenic growth responses in detail. Using *in vitro* kinase assays, we first demonstrate that AFC1, AFC2, and AFC3 show distinct temperature-activity profiles within the physiological temperature range of Arabidopsis that depend on the conserved activation segment and the unstructured N-terminal region of the protein. We then used CRISPR/Cas9 to generate single and higher order *afc* mutants to resolve the functional role of AFCs in thermomorphogenic growth responses and signalling. Our data show that AFCs are required for thermoresponsive hypocotyl elongation as well as the establishment of temperature-dependent splicing patterns. Finally, we identify SR34 and SR34a as phosphorylation targets that mediate thermomorphogenic hypocotyl elongation growth downstream of AFC activity. Therefore, we propose that AFC kinases translate ambient temperature information into a cellular signal through phosphorylation of specific SR proteins, thereby controlling warm temperature-dependent alternative splicing and thermoresponsive growth in Arabidopsis.

## Results

### AFC1, AFC2 and AFC3 show distinct temperature-activity profiles mediated by the unstructured N-terminus and the activation segment

Our recent work showed that animal CLKs display highly temperature-dependent activities with an off-switch at the upper limit of the physiologically relevant temperature range in various endothermic species ^23^. Based on this finding, we aimed at investigating whether the activities of the homologous Arabidopsis kinases, AFC1 (At3g53570), AFC2 (At4g24740), and AFC3 (At4g32660) (Supplementary Fig. 1a), are also temperature-dependent. A comparison of the predicted three-dimensional structures of the three AFC kinase domains with the experimentally resolved structure of human CLK1 yielded a high degree of structural homology with all major sequence elements being conserved and only minor differences, mainly in unstructured regions (Fig. 1a). Subsequent *in vitro* kinase assays revealed a prominent temperature sensitivity of all three kinases with distinct temperature-activity profiles. While AFC1 activity was highest at lower temperatures between 4 °C and 20 °C, AFC3 had its highest activity between 24 °C and 32 °C (Fig. 1b, c; Supplementary Fig. 1b). In contrast, AFC2 showed largely stable activity between 4 °C and 28 °C. Notably, the *in vitro* activity of all three kinases declined rapidly at temperatures above 32 °C.

**Figure 1.**
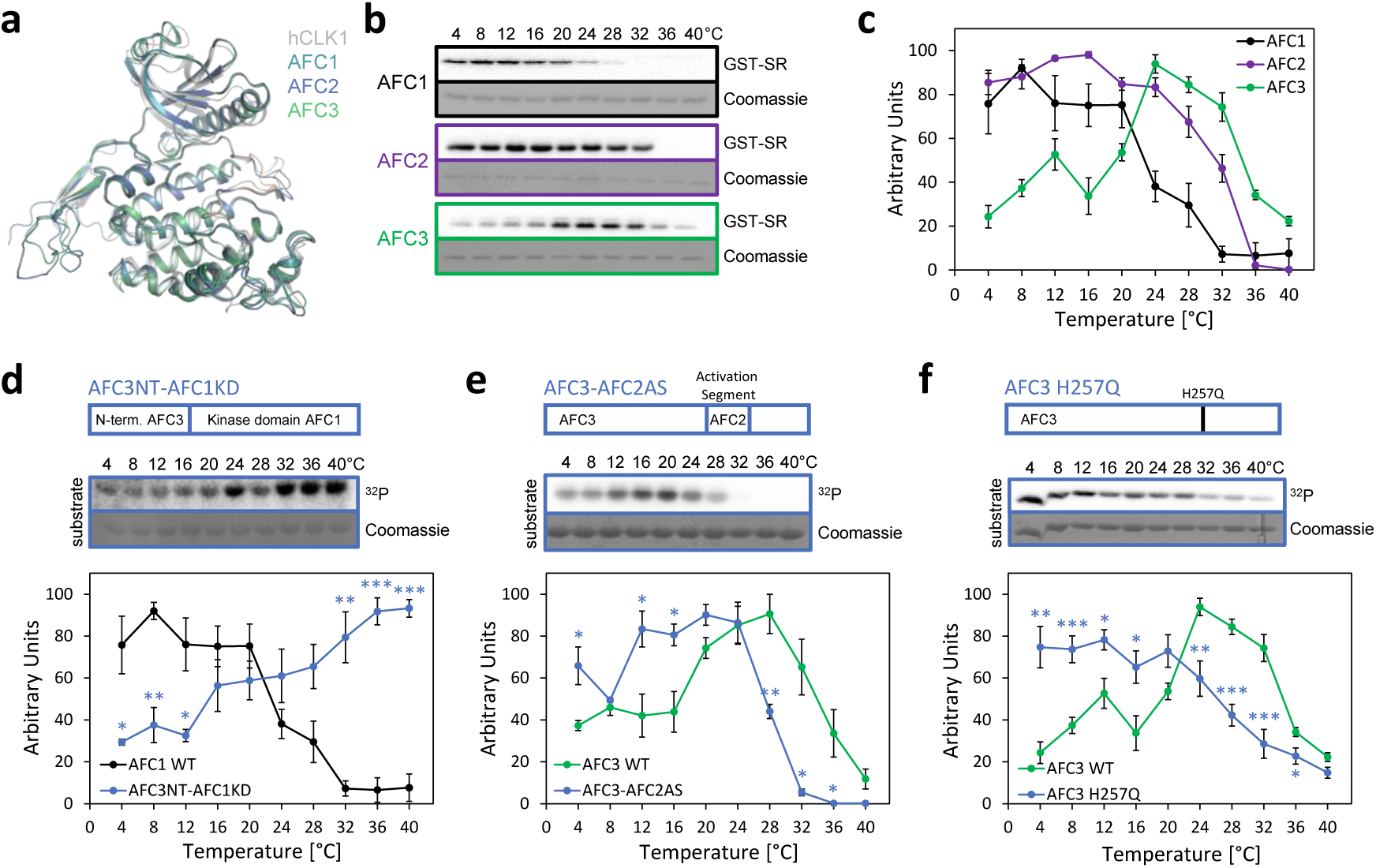
AFC1, AFC2 and AFC3 show distinct temperature-activity profiles mediated by the unstructured N-terminus and the activation segment. **(a)** Superimposition of the predicted kinase domains of AFC1 (P51566, residues 102-467), AFC2 (P51567, residues 85-427) and AFC3 (P51568, residues 58-400) with the crystal structure of human CLK1 kinase domain (PDB: 6TW2). **(b)** Temperature-dependent phosphorylation of a GST-SR substrate by AFC1, AFC2 and AFC3. After separation by SDS-PAGE, phosphorylation of synthetic GST-SR was detected by autoradiography (top, ^32^P). Equal loading was confirmed by Coomassie staining (bottom). Loading controls for the three kinases are shown in Supplementary Fig. 1b. **(c)** Quantification of temperature-dependent substrate phosphorylation activity of AFC1, AFC2 and AFC3. For quantification, the highest signal intensity within an assay was set to 100 and used for normalisation. Shown are mean values ± SE; *n* = 3 for AFC1 and *n* = 6 for AFC2 and AFC3, respectively. **(d-f)** Temperature-dependent activity of chimeric AFC1 containing the AFC3 N-terminus (d), chimeric AFC3 containing the AFC2 activation segment (e), and an AFC3 H257Q point mutant (f). Reaction temperatures are indicated on top. In all cases, representative images of synthetic GST-SR phosphorylation (autoradiography, ^32^P, top) and the respective substrate loading controls (Coomassie staining, bottom) are shown. Displayed are mean values ± SE from *n* = 3 (d), *n* = 3 (e), and *n* = 6-7 (f) replicates. Statistically significant differences were determined by Student’s *t*-tests (*: *p* < 0.05, **: *p* < 0.01, ***: *p* < 0.001). Note that data for AFC1 depicted in (d) and AFC3 depicted in (f) are the same as presented in (c). WT: wild type.

To address the molecular details underlying the opposing temperature-activity profiles of AFC1 and AFC3, we created several chimeric proteins and mutants focusing on the unstructured N-terminus and specific amino acids within the activation segment, which are known to set the specific temperature-activity profile of animal CLKs^23^. We first generated a chimeric kinase consisting of the AFC3 N-terminus and the AFC1 kinase domain. This chimera showed a temperature-activity profile rather resembling that of AFC3 than that of AFC1, showing that the N-terminus is involved in setting the temperature-controlled activity. In addition, the chimeric kinase remained active at temperatures above 28 °C (Fig. 1d), suggesting that the N-terminus plays a role in stabilising the kinase domain or the active centre. Furthermore, this stabilising effect seems to depend on the exact sequence context, as it was absent in full-length AFC3. The kinase domains alone displayed an activity similar to the full-length kinases (Supplementary Fig. 1c), showing that the altered activity of the AFC1-AFC3 chimera was mediated by the presence of the AFC3 N-terminus and not the absence of the AFC1 N-terminus.

We then created a chimeric isoform of AFC3 containing the AFC2 activation segment (Fig. 1e). This chimera showed a temperature-activity profile reminiscent of the behaviour of both wild-type (WT) kinases, as the activity remained higher in the temperature range between 12 °C and 24 °C, as seen in AFC2, and decreased at lower temperatures, as seen in AFC3.

Finally, two amino acid residues, R343 and H344, are instrumental in mediating the temperature sensitivity of animal CLK1, as they change their conformation in a temperature-dependent manner potentially blocking substrate access to the active centre at higher temperature^23^. These amino acids are conserved in all three AFCs (Supplementary Fig. 1a). We thus tested whether they were also involved in the temperature-dependent regulation of AFC activity. While an AFC1 H304Q mutation had little effect on the temperature-activity profile (Supplementary Fig. 1d), the corresponding AFC2 H285Q (Supplementary Fig. 1e) and AFC3 H257Q (Fig. 1f) mutants showed strongly altered temperature-activity profiles compared to the respective WT isoforms. This suggests that a mechanism similar to the one that regulates temperature-controlled activity of CLKs also controls AFC2 and AFC3 activity. Altogether, these data establish AFC activity as highly temperature-responsive in the range between 4 °C and 40 °C with striking differences between AFC1, AFC2, and AFC3. Furthermore, we show that at least two protein domains, the unstructured N-terminus and the activation segment, are involved in mediating temperature sensitivity and that the temperature-activity profile can be modulated by mutating specific amino acids.

### A glutamine centred H-bond network mediates AFC3 activity at high temperatures

To further understand the molecular and structural basis underlying the different temperature-activity profiles of AFC1 and AFC3, we examined the predicted structures in more detail. Superposition of the AFC1 and AFC3 kinase domains revealed a high similarity with a root mean square deviation (RMSD) of 1.25 Å and a TM-score of 0.92 with 69% identity (Fig. 2a). Structural differences were restricted to a flexible loop in the C lobe and the also flexible, but catalytically relevant activation segment.

**Figure 2.**
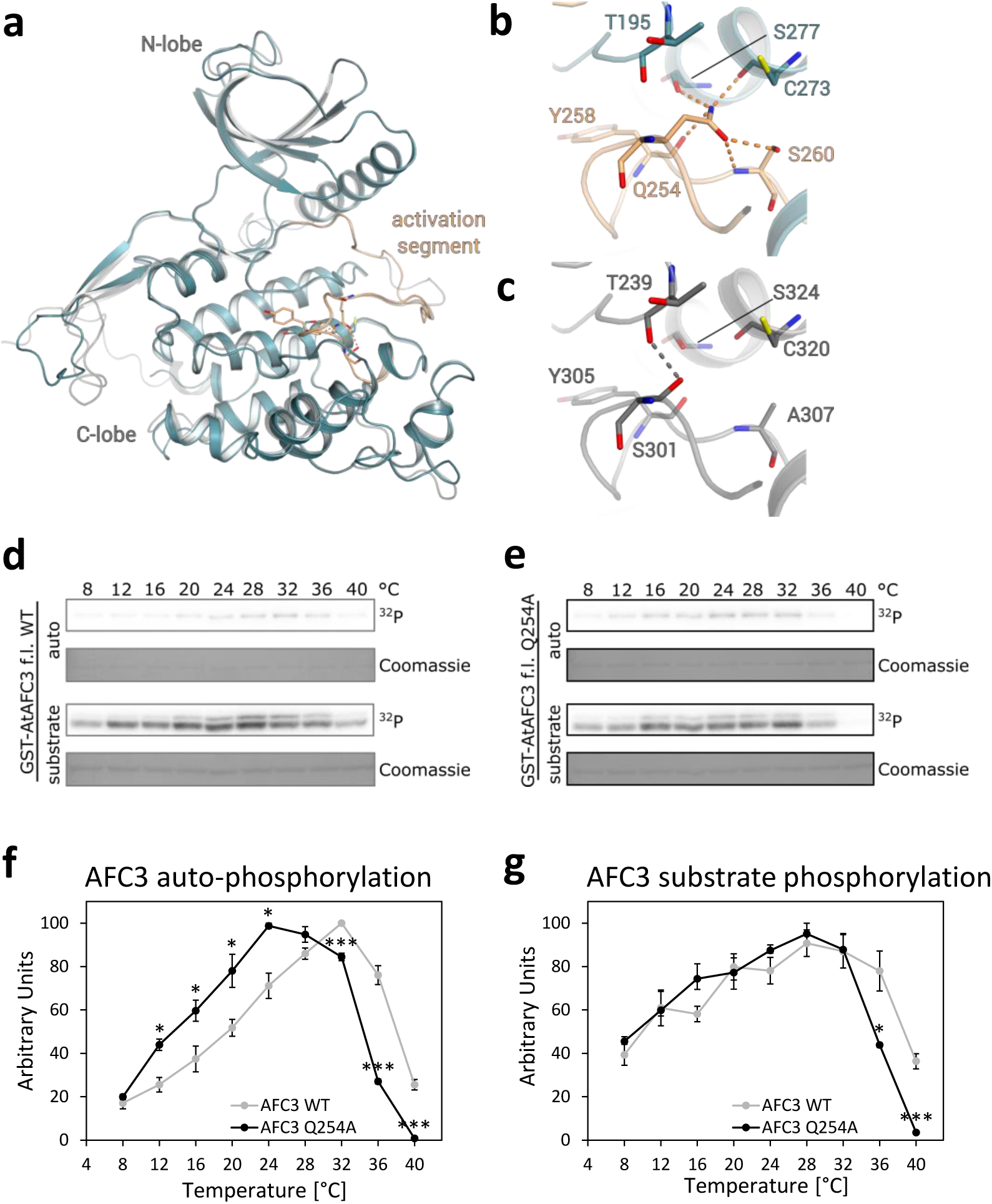
A glutamine centred H-bond network mediates AFC3 activity at high temperatures. **(a)** Superimposition of the predicted structures of the kinase domains of AFC1 (P51566, residues 102-467, grey) and AFC3 (P51568, residues 58-400, teal) with the AFC3 activation segment coloured in light orange. Residues of AFC3 involved in the formation of the H-bond network around AFC3 Q254 are shown as stick models. **(b,c)** Enlarged view of the H-bond network around AFC3 Q254 (b) and AFC1 S301 (c). Interacting residues of AFC3 and the corresponding AFC1 residues are depicted as sticks with the formed H-bonds (distance cut-off = 3.2 Å) coloured in orange (AFC3) or grey (AFC1). **(d,e)** Representative images of temperature-dependent auto- and substrate phosphorylation of AFC3 WT (d) and an AFC3 Q254A mutant (e). Phosphorylation of GST-tagged AFCs and synthetic GST-SR was detected by autoradiography (top, ^32^P). Equal loading was confirmed by Coomassie staining (bottom). **(f,g)** Quantification of temperature-dependent auto-phosphorylation (f) and substrate phosphorylation (g) activity of AFC3 WT and AFC3 Q254H mutant. For quantification, the highest signal intensity within an assay was set to 100 and used for normalisation. Shown are mean values ± SE from *n* = 4 (AFC3 WT) and *n* = 3 (AFC3 Q254A) replicates. Statistically significant differences were determined by Student’s *t*-tests (*: *p* < 0.05, **: *p* < 0.01, ***: *p* < 0.001).

For the activation segment of AFC3, we observed an H-bond network involving Q254 and the sidechains of S277 from the αF-helix and S260, which is also part of the activation segment, and the backbones of C273, S260 and Y258 (Fig. 2b). These interactions may stabilise the region of the activation segment containing R256 and H257 in an active conformation. As discussed above, the homologous residues in human CLK1 are essential for defining the temperature-activity optimum^23^ and an H257Q mutation also resulted in a strongly altered temperature-activity profile of AFC3 (Fig. 1f). In AFC1, which is not active at temperatures above 32 °C (Fig. 1b, c), the amino acids corresponding to AFC3 Q254 and S260 are S301 and A307, respectively. Since S301 only forms a single H-bond in AFC1, this part of the activation segment lacks stabilising interactions (Fig. 2c). Thus, we hypothesised, that the H-bond network around AFC3 Q254 stabilises R256 and H257 in an active conformation at higher temperatures, thereby allowing AFC3 activity at temperatures where AFC1 is already inactive. To test this hypothesis, we generated an AFC3 Q254A mutant and performed *in vitro* kinase assays (Fig. 2d, e). Indeed, temperature-responsive autophosphorylation activity was shifted towards lower temperatures by several degrees for the AFC3 Q254A mutant compared to WT AFC3, including a substantial decline in activity at temperatures above 32 °C (Fig. 2f). Concerning substrate phosphorylation, no significant differences were observed between both variants between 4 °C and 32 °C (Fig. 2g). However, the Q254A mutation led to a more pronounced decrease in substrate phosphorylation above 32 °C than was observed for WT AFC3, yielding a similar effect as for autophosphorylation. In conclusion, these data show that residue Q254 is required for the activity of AFC3 at temperatures above 32 °C.

### AFC1 and AFC3 are required for thermoresponsive growth

Based on the observed temperature-sensitive activity of the three AFC kinases, we hypothesised that they could be involved in the regulation of temperature-dependent growth responses *in vivo*. Initial support for this hypothesis came from the observation that warm temperature-dependent hypocotyl elongation in eight-day-old Arabidopsis Columbia-0 (Col-0) seedlings was negatively affected by the CLK1/4 inhibitor TG003^30^, which also inhibited AFC activity, in a dose-dependent manner (Fig. 3a; Supplementary Fig. 2a-d). We then generated *afc* single and higher order mutant lines using CRISPR/Cas9 (Supplementary Fig. 3a-c; Supplementary Fig. 4a-c). The *afc* single mutants did not show any differences in temperature-dependent hypocotyl elongation (Fig. 3b). However, hypocotyl growth was significantly reduced in warm temperature-exposed seedlings of two independent *afc1 afc2 afc3* triple mutant lines designated as *afc1/2/3 #1* and *afc1/2/3 #2* (Fig. 3c, d). At the same time no difference in hypocotyl length between the two mutant lines and the WT was observed at 17 °C and 21 °C. In addition, we observed a strongly reduced thermonastic leaf movement response in *afc* triple mutant plants (Supplementary Fig. 5a-c) as well as a modest reduction in warm temperature-dependent petiole elongation (Supplementary Fig. 5d), while overall vegetative growth was not affected in *afc1/2/3 #1* and *#2* (Supplementary Fig. 5e). Complementation of the *afc1/2/3 #2* mutant with the genomic sequence of *AFC3* under its native promoter led to a partial restoration of warm temperature-dependent hypocotyl elongation (Fig. 3e, f). Skotomorphogenic elongation growth was not affected in the two *afc1/2/3* triple mutant lines (Supplementary Fig. 5f, g) indicating their capacity for WT-like elongation growth. Moreover, *afc1/2/3 #2* did not show any differences in short- or long-term thermotolerance compared to WT when exposed to heat stress above 37 °C (Supplementary Fig. 5h, i). Finally, the temperature-dependent hypocotyl elongation of *afc1/2* and *afc1/3* double mutants was assessed to reveal potential functional redundancies between the three AFC kinases during thermomorphogenesis. While *afc1/2* seedlings displayed an intermediate elongation phenotype (Fig. 3g) thereby resembling *AFC3*-complemented *afc1/2/3* triple mutant seedlings (Fig. 3e), *afc1/3* was indistinguishable from *afc1/2/3 #2* (Fig. 3g) suggesting that AFC2 cannot compensate for a loss of AFC1 and AFC3 during thermomorphogenesis. While our data indicate that AFC1 and AFC3 are specifically required for warm temperature-dependent growth responses in Arabidopsis, we continued molecular analyses with the triple mutant plants to avoid potential confounding effects.

**Figure 3.**
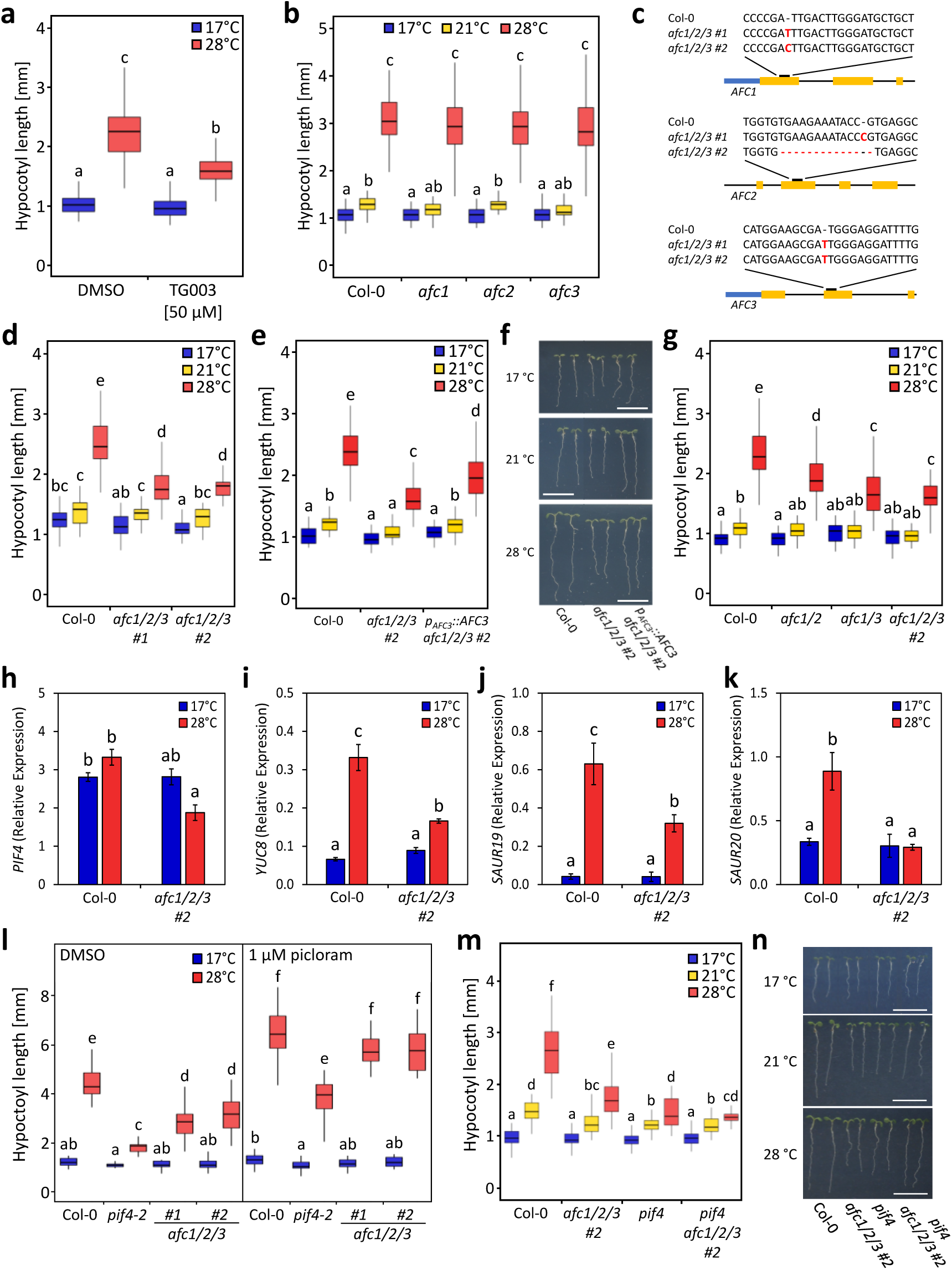
AFCs are required for thermoresponsive growth. **(a)** Hypocotyl lengths of eight-day-old Col-0 seedlings supplemented with the AFC kinase inhibitor TG003 and exposed to 17 °C or 28 °C. Seedlings were grown on non-supplemented medium for three days and then transferred to TG003- or DMSO-containing (solvent control) medium. On day five half of the plates were shifted to 28 °C while the other half remained at 17 °C (*n* = 39-48 seedlings per treatment and temperature). Inhibition of AFC activity by TG003 was confirmed *in vitro* for recombinant AFC3 (Supplementary Fig. 2a). **(b)** Hypocotyl lengths of seven-day-old Col-0, *afc1*, *afc2* and *afc3* single mutant seedlings at 17 °C, 21 °C and 28 °C. Seedlings were grown at 17°C for four days and then exposed to the indicated temperatures for three days (*n* = 43-59 seedlings per genotype and temperature). **(c)** Overview of the genetic modifications in the two independent *afc1 afc2 afc3* triple mutant lines generated by CRISPR/Cas9 and designated as *afc1/2/3 #1* and *afc1/2/3 #2*. **(d)** Hypocotyl lengths of seven-day-old seedlings of Col-0, *afc1/2/3 #1* and *afc1/2/3 #2* at 17 °C, 21 °C and 28 °C (*n* = 29-40 seedlings per genotype and temperature). **(e,f),** Hypocotyl lengths (e) and representative images (f) of seven-day-old seedlings of Col-0, *afc1/2/3 #2* and *afc1/2/3 #2* complemented with genomic *AFC3* under control of the *AFC3* promotor (*p_AFC3_::AFC3 afc1/2/3 #2*) at 17 °C, 21 °C and 28 °C (*n* = 31-57 seedlings per genotype and temperature). **(g)** Hypocotyl lengths of seven-day-old seedlings of Col-0, *afc1/2*, *afc1/3* and *afc1/2/3 #2* at 17 °C, 21 °C and 28 °C (*n* = 25-47 seedlings per genotype and temperature). **(h-k)** Relative expression of *PIF4* (h), *YUCCA8* (i), *SAUR19* (j) and *SAUR20* (k) in seven-day old Col-0 and *afc1/2/3 #2* seedlings as determined by RT-qPCR (mean ± SE, *n* = 3-4). Seedlings were grown at 17 °C for four days and exposed to 28 °C for another three days while control seedlings were kept at 17 °C throughout the experiment. **(l)** Hypocotyl lengths of seven-day-old Col-0, *pif4-2*, *afc1/2/3 #1* and *#2* seedlings supplemented with 1 µM of the synthetic auxin picloram and exposed to 17 °C or 28 °C for three days (*n* = 16-32 seedlings per treatment and temperature). **(m,n)** Hypocotyl lengths (m) and representative images (n) of seven-day-old seedlings of Col-0, *afc1/2/3 #*2, *pif4* and a *pif4 afc1/2/3 #2* quadruple mutant at 17 °C, 21 °C and 28 °C (*n* = 31-57 seedlings per genotype and temperature). Seedlings were grown at 17 °C for four days and then exposed to the indicated temperatures for three days. Statistically significant differences were determined by two-or three-way ANOVAs followed by Tukey HSD *post-hoc* tests and are indicated by different letters above boxes (*p* < 0.05). Scale bars represent 1 cm.

### AFCs interfere with PIF4-dependent signalling

Given the central role of PIF4 in thermomorphogenesis signalling, we next assessed whether the expression of *PIF4*-regulated genes was affected in *afc1/2/3* triple mutant seedlings. While *PIF4* itself did not show any significant transcriptional regulation in response to warm temperature irrespectively of plant genotype, induction of the PIF4-dependent genes *YUCCA8* (*YUC8*), *SMALL AUXIN UP RNA 19* (*SAUR19*) and *SAUR20* was significantly reduced in *afc1/2/3 #2* compared to the WT following exposure to 28 °C (Fig. 3h-k). Enzymatic YUC8 is required for warm temperature-induced auxin biosynthesis thereby mediating hypocotyl elongation^31^. We thus tested whether pharmacological supplementation with the synthetic auxin picloram could restore warm temperature-dependent hypocotyl elongation growth in *afc1/2/3*. When supplemented with picloram, *afc1/2/3 #1* and *afc1/2/3 #2* displayed WT-like hypocotyl lengths following exposure to 28 °C and hypocotyl elongation growth in the *pif4-2* mutant^32^ was also partially rescued (Fig. 3l). Notably, in the absence of picloram elongation growth was more strongly reduced in *pif4-2* compared to the two *afc1/2/3* triple mutant lines at 28 °C (Fig. 3l). Finally, we generated a *pif4 afc1/2/3 #2* quadruple mutant using CRISPR/Cas9 to determine whether AFCs function in the same pathway as PIF4 (Supplementary Fig. 6a-d). Elongation growth of the *pif4 afc1/2/3 #2* quadruple mutant was indistinguishable from that of the *pif4* single mutant carrying the same mutant allele but in WT background (Fig. 3m, n). In agreement with the previous experiment, the hypocotyls of *afc1/2/3 #2* were slightly longer at 28 °C compared with the *pif4* single mutant. In conclusion, our data suggest that AFCs function in the same pathway as PIF4 during warm temperature-induced hypocotyl elongation.

### AFCs regulate temperature-dependent splicing of genes involved in RNA processing

Next, we aimed at investigating the temperature-dependent splicing patterns in the *afc* triple mutant lines in more detail. To this end, we performed RNA-sequencing (RNA-seq) on seven-day old *afc1/2/3 #1* and *afc1/2/3 #2* seedlings following exposure to 28 °C for 8 and 24 hours, respectively, and compared their splicing patterns to those of WT seedlings (Fig. 4a). Subsequent principal component analysis on all detected splice events yielded a clear separation between the different genotypes and temperature conditions indicating a genotype-dependent effect of temperature on the global splicing patterns (Fig. 4b). We then assessed whether AFCs are involved in the regulation of a specific type of alternative splicing in response to a temperature increase. Therefore, we first identified genes that were differentially alternatively spliced (DAS) after the temperature shift in WT and triple mutant lines by pairwise comparisons and subsequently analysed the relative distribution of the four types of alternative splicing. However, we did not observe any alterations concerning the frequencies of the different alternative splicing types between the two mutant lines and the WT. Irrespectively of plant genotype, and expected based on previous observations, intron retention (IR) was the dominant type of temperature-dependent alternative splicing, followed by exon skipping (ES), and alternative usage of 3’ (Alt 3’) and 5’ (Alt 5’) splice sites (Fig. 4c).

**Figure 4.**
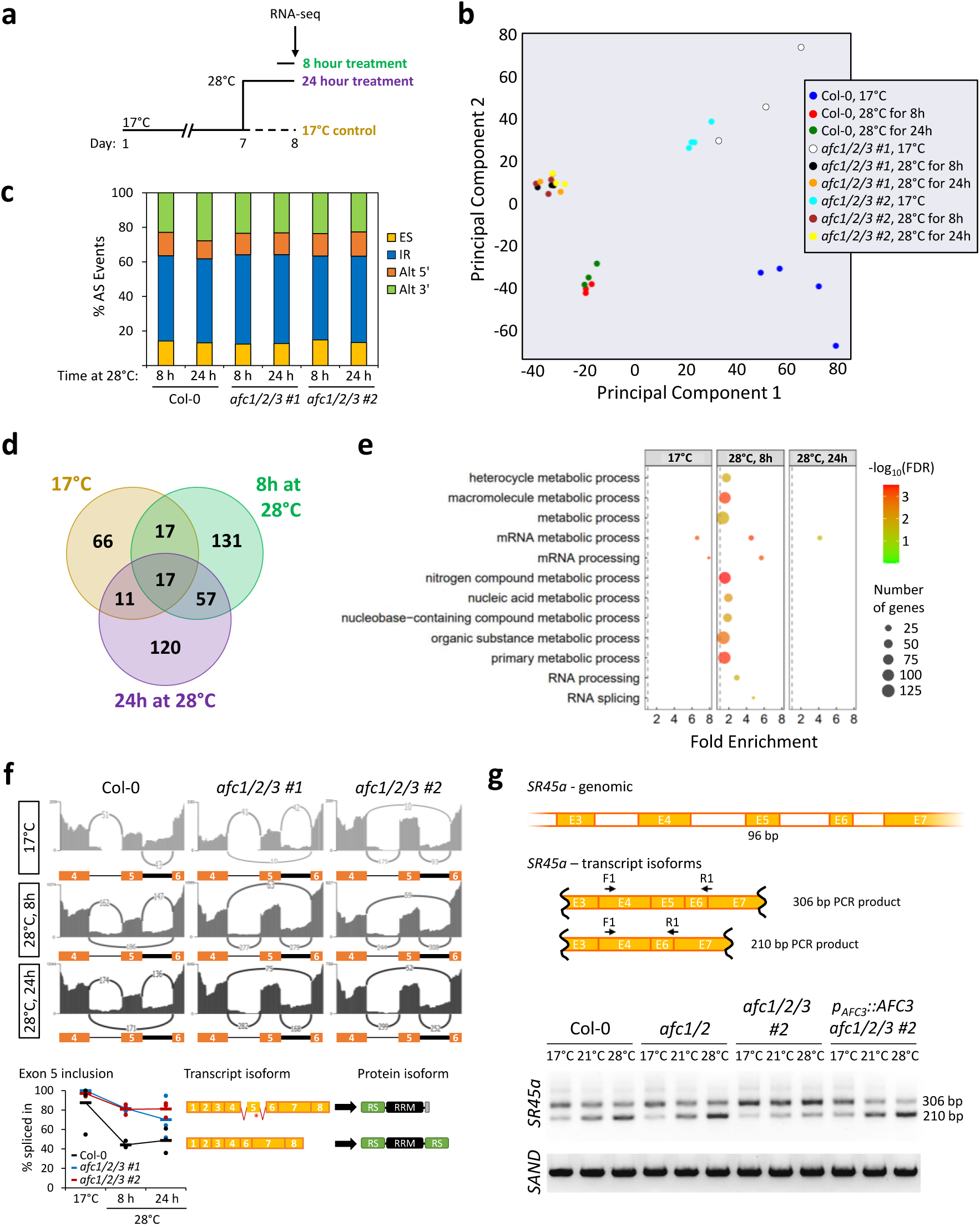
AFCs regulate the temperature-dependent splicing of genes that are involved in RNA processing. **(a)** Schematic overview of the experimental conditions used for the analysis of temperature-dependent splicing patterns. Seedlings of Col-0, *afc1/2/3 #1* and *afc1/2/3 #2* were grown at 17 °C and then exposed to 28 °C for 24 hours starting at the end of day 7 or exposed to 28 °C for 8 hours starting on day 8. Control seedlings remained at 17 °C. All samples were taken at Zeitgeber Time (ZT) 11 of day 8, i.e., 11 hours after the onset of light. **(b)** Principal component analysis of all detected splice events. The different sample groups are indicated by different colours. **(c)** Relative frequencies of the different alternative splicing types among the differentially alternatively spliced genes in Col-0, *afc1/2/3 #1* and *afc1/2/3 #2* following exposure to 28 °C for 8 or 24 hours. **(d)** Intersection of genes that were differentially alternatively spliced in both *afc1/2/3 #1* and *afc1/2/3 #2* compared to Col-0 at the different temperature conditions. **(e)** Gene ontology term enrichment analysis for genes that were DAS in both *afc1/2/3 #1* and *afc1/2/3 #2* compared to Col-0 at the different temperature conditions. **(f)** Temperature-dependent splicing of *SR45a* exon 5 in Col-0, *afc1/2/3 #1* and *afc1/2/3 #2* as determined by RNA-sequencing. Representative Shashimi plots for the indicated temperatures (top) and the rMATS-derived quantification of exon 5-inclusion (bottom) are shown with the functional consequence of exon 5-inclusion indicated on the right (*n* = 3-4). **(g)** Temperature-dependent splicing of *SR45a* exon 5 in Col-0, *afc1/2*, *afc1/2/3 #2* and *afc1/2/3 #2* complemented with genomic *AFC3 (p_AFC3_::AFC3 afc1/2/3 #2*) as determined by reverse transcription PCR. A schematic overview of the *SR45a* genomic region and the respective splice products are shown on top with the primers used indicated above and the expected PCR product sizes on the right. Seedlings were grown at 17 °C for four days and then exposed to the indicated temperatures for another three days. Amplification of *SAND* was used as expression control. Additional controls and replicates are shown in Supplementary Fig. 6.

Next, we were interested in the genes that showed genotype-dependent differences in their splicing patterns at the different temperatures (Fig. 4d; Supplementary Data 1). Considering only genes that were affected in both mutant lines compared to WT yielded 111 differentially alternatively spliced genes (DASGs) under control conditions (Supplementary Data 2a). At 28 °C, 222 and 205 DASGs were detected after 8 and 24 hours, respectively, suggesting a higher impact of AFCs on alternative splicing at warm temperatures. Out of the 308 unique genes that were DAS in *afc1/2/3 #1* and *afc1/2/3 #2* at 28 °C, 41% were spliced in a temperature-dependent manner in the WT, including several genes coding for SR proteins (Supplementary Data 2b). Subsequent gene ontology (GO) term enrichment analysis of the DASGs likewise yielded a significant enrichment of several terms associated with RNA metabolism and processing, especially at the early time point of the 28 °C treatment (Fig. 4e; Supplementary Data 3).

Since the alternative splicing of SR protein genes may result in altered SR protein expression and/or activity and thus be instrumental in the establishment of global temperature-dependent alternative splicing patterns, we examined the impact of AFC activity on the temperature-dependent splicing of SR protein genes and the resulting consequences in more detail. As one example we chose the non-classical SR protein SR45a (At1g07350). Under control conditions in WT nearly all detected transcripts of *SR45a* included exon 5, which contains a premature termination codon (PTC) and gives rise to a truncated protein isoform that lacks the C-terminal RS domain (Fig. 4f). Exposure to 28 °C, however, induced skipping of exon 5 in ∼50% of the detected *SR45a* transcripts in WT but not in *afc1/2/3 #1* or *afc1/2/3 #2* seedlings. Exon 5-skipping produces the transcript isoform that encodes full-length SR45a containing the C-terminal RS domain. These findings were confirmed through reverse transcription-PCR in an independent experiment, which again showed that the *afc1/2* double mutant behaved like the WT (Fig. 4g; Supplementary Fig. 7a-d). Moreover, complementation with genomic *AFC3* rescued the defective temperature-dependent splicing of *SR45a* in *afc1/2/3 #2* (Fig. 4g). Thus, the warm temperature-induced shift from truncated to full-length SR45a in WT seedlings is dependent on AFC kinases.

In another example, we observed genotype-specific differences in the temperature-regulated splicing of *RS40* (At4g25500). In WT seedlings, exposure to elevated temperature elicited an increase in the abundance of a nonsense-mediated mRNA decay-sensitive *RS40* transcript isoform via inclusion of exon 4, which correlated with a decrease in total *RS40* transcript levels (Supplementary Fig. 7e). By contrast, inclusion of PTC-containing exon 4 was already enhanced in both *afc1/2/3 #1* and *afc1/2/3 #2* under control conditions and increased even further at 28 °C. This also resulted in lower total *RS40* transcript levels, suggesting AFC-dependent post-transcriptional regulation of *RS40* expression, which is likely further controlled through retention of intron 3 (Supplementary Fig. 7e). In conclusion, our data provide evidence that AFCs are involved in the regulation of temperature-dependent alternative splicing and that they particularly control the correct splicing of a subset of genes that are involved in RNA processing themselves.

### SR proteins contribute to temperature-induced hypocotyl elongation

Since SR proteins serve as potential targets of AFC kinases^25, 33^, we next aimed at investigating their role during seedling thermomorphogenesis in more detail. To this end, we first investigated the temperature-dependent expression of all 20 genes coding for SR proteins using our RNA-seq data (Supplementary Data 4). While only two *SR* genes were significantly downregulated after exposure to 28 °C in WT seedlings, six *SR* genes were significantly upregulated (|log_2_FC| > 1, *p* < 0.05) (Fig. 5a; Supplementary Data 4) confirming the previously reported temperature-dependent regulation of SR protein expression^19^. Moreover, seven *SR* genes were differentially expressed (DE) in *afc1/2/3 #1* and *afc1/2/3 #2* compared to the WT with six of them showing reduced transcript levels in the mutant seedlings (Fig. 5a; Supplementary Data 4). Notably, there was little overlap between *SR* genes that were DE and DAS in *afc* triple mutants compared to the WT, with the two cases showing DE and DAS likely reflecting regulation by alternative splicing coupled to nonsense-mediated mRNA decay (AS-NMD). It is also worth noting that 12 out of 20 *SR* genes showed DE and/or DAS in the *afc* triple mutant, representing more than 50% of *SR* genes, which very likely contributes to global DAS in the mutant plants.

**Figure 5.**
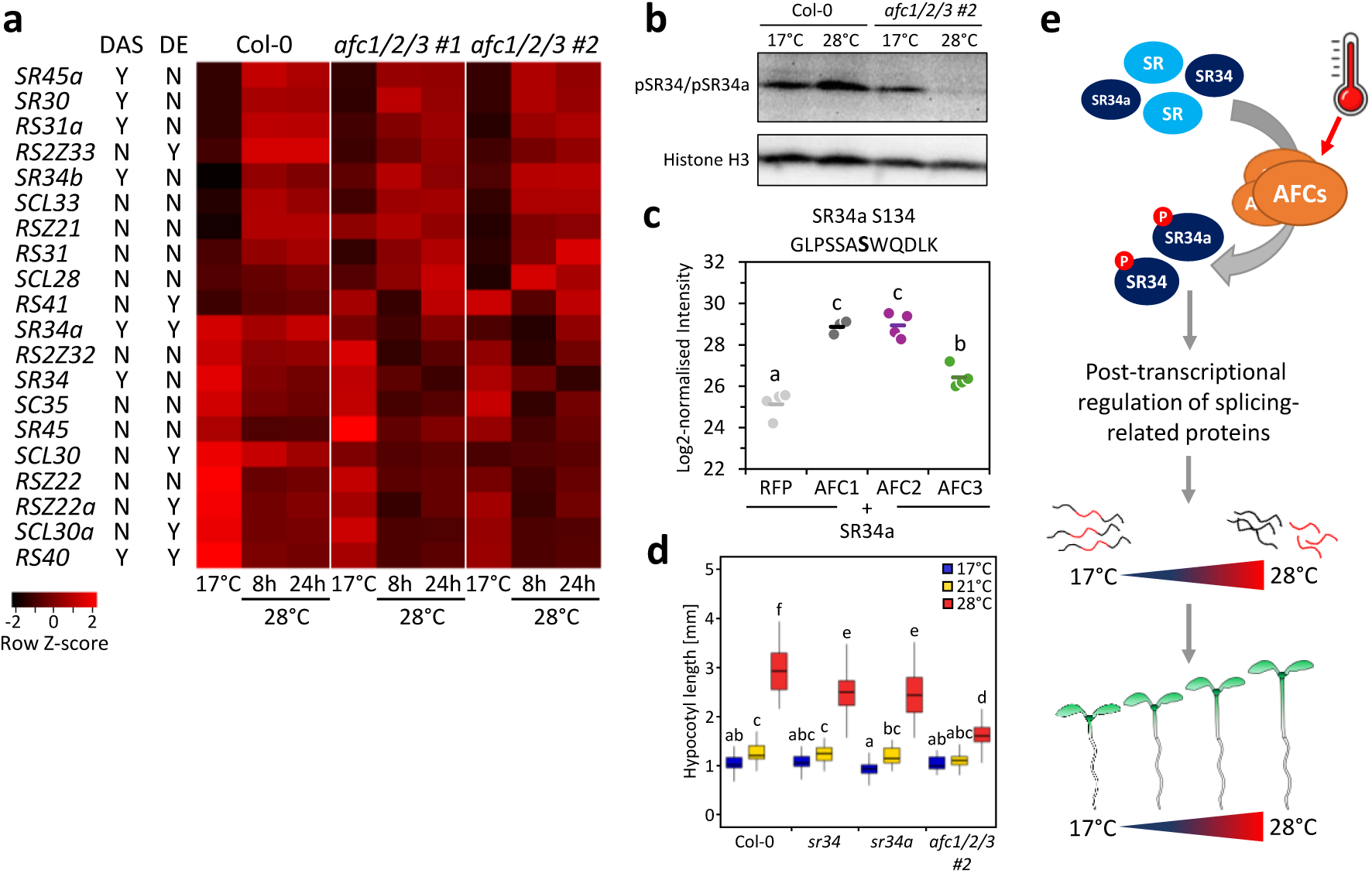
SR proteins contribute to warm temperature-dependent hypocotyl elongation. **(a)** Temperature-dependent expression of SR proteins in seven-day-old seedlings of Col-0, *afc1/2/3 #1* and *afc1/2/3 #2*. Seedlings were exposed to 28 °C for the indicated time periods and transcript levels determined by RNA-sequencing. Genes that were differentially expressed (DE) or differentially alternatively spliced (DAS) in both *afc* triple mutants compared to Col-0 under at least one of the three temperature conditions are indicated next to the gene name. Y: yes, N: no. **(b)** Analysis of SR protein phosphorylation status in nine-day-old seedlings of Col-0 and *afc1/2/3 #2* that were exposed to 28 °C for one hour or remained at 17 °C as control. Phosphorylation was assayed using an α-pan-phosphoepitope SR-specific antibody and an α-histone H3 antibody for normalisation. **(c)** Log2 intensities of an SR34a phosphopetide containing Ser134 (in bold) as detected by mass spectrometry following co-expression of SR34a-GFP with RFP-tagged AFC1, AFC2 or AFC3 in *N. benthamiana* leaves. Horizontal bars indicate mean values (*n* = 4). **(d)** Hypocotyl lengths of seven-day-old seedlings of Col-0, *sr34*, *sr34a* and *afc1/2/3 #2* at 17 °C, 21 °C and 28 °C. Seedlings were grown at 17 °C for four days and then exposed to the indicated temperatures for three days (*n* = 25-43 seedlings per genotype and temperature). Statistically significant differences were determined by two-way ANOVA followed by a Tukey HSD *post-hoc* test and are indicated by different letters above boxes (*p* < 0.05). **(e)** Current working model on the function of AFC kinases during thermomorphogenic growth responses. Upon exposure to a temperature increase, AFCs phosphorylate specific SR proteins, likely SR34 and SR34a. This in turn leads to the post-transcriptional regulation of splicing-related genes including genes coding for SR proteins, thereby contributing to the establishment of global warm temperature-dependent alternative splicing patterns. Correct splicing at elevated temperatures is required for proper thermomorphogenic growth responses. Consistently, *afc* triple mutant seedlings show altered temperature-dependent alternative splicing patterns and reduced hypocotyl elongation growth at 28 °C.

Given frequent auto- and cross-regulation of SR proteins^19, 34, 35, 36^, altering the activity of only one SR protein by phosphorylation could induce changes in DAS and DE of additional *SR* genes, which could then mediate more global effects. To characterize SR protein candidates that are potentially phosphorylated by AFC kinases in a temperature-dependent manner, we identified SR proteins that are rapidly phosphorylated in WT seedlings after exposure to 28 °C in two datasets, a first one from Arabidopsis seedlings exposed to elevated temperature during the night^22^ (Supplementary Data 5a) and a second one from Arabidopsis seedlings exposed to elevated temperature during the day (Supplementary Data 5b). Comparing these two datasets revealed common regulation of SR34, SR34a and SCL30, but only for SR34 (S273) and SR34a (S134) the same phosphosites were detected in both datasets (Supplementary Fig. 8a) making these good candidates for further investigation. To establish SR34 and SR34a as targets of warm temperature-dependent AFC activity, we assessed SR protein phosphorylation in wild-type and *afc1/2/3 #2* seedlings using an α-pan-phosphoepitope SR-specific antibody^23^. In agreement with temperature-dependent SR34 and SR34a phosphorylation, we observed an increased signal intensity at approximately 32 kDa in wild-type seedlings that were exposed to 28 °C for one hour (Fig. 5b; Supplementary Fig. 8b-f). By contrast, signal intensity was strongly reduced in warm temperature-treated *afc* triple mutant seedlings compared to the 17 °C control. Transient co-expression of GFP-tagged SR34a together with AFC1, AFC2 or AFC3 in *N. benthamiana* leaves followed by immunoprecipitation-mass spectrometry confirmed the AFC-dependent phosphorylation of SR34a at S134 (Fig. 5c), the site that was also phosphorylated in a temperature-dependent manner in WT. In addition to S134, we detected several other SR34a phosphosites that were phosphorylated in an AFC-dependent manner (Supplementary Data 5c). Moreover, phosphopeptides for all three AFCs were detected following immunoprecipitation of GFP-tagged SR34a suggesting an interaction between SR34a and the three AFC kinases (Supplementary Fig. 8g-i; Supplementary Data 5d). Finally, we assessed thermoresponsive hypocotyl elongation of *sr34* and *sr34a* knockdown mutant seedlings (Supplementary Fig. 9a, b) and observed a significant reduction for both genotypes (Fig. 5d). Taken together, these data indicate a partially redundant functional role of SR34 and SR34a in seedling thermomorphogenesis downstream of AFC activity.

## Discussion

Plants are able to actively adjust their growth and development to the prevalent temperature conditions^2^. This, however, requires the accurate perception and integration of temperature information with cellular processes and signalling^37^. Here, we have shown that AFC kinases integrate temperature information with the splicing machinery to control temperature-dependent alternative splicing and thermomorphogenic growth responses.

Thermosensory properties had been demonstrated previously for the human homologs CLK1 and CLK4^23^. In contrast to their mammalian counterparts, the three plant AFCs showed distinct temperature-activity profiles indicating a functional diversification that is likely adapted to the broader temperature range plants are exposed to, and which contrasts with the tightly controlled core body temperature of endotherms^23^. This hypothesised temperature adaptation is further supported by the observation that CLKs from poikilothermic animals change their activity within the physiological temperature range of the respective host organism^23^.

From a mechanistic point of view the structural elements that determine the temperature-responsive activity of animal CLKs and plant AFCs are identical, including the unstructured N-terminus and the activation segment. However, our data suggest that additional, so far unknown, mechanisms contribute to AFC temperature sensitivity. This is supported by the observation that the targeted mutation of H257 within the RH motif of AFC3 led to a shift of maximum kinase activity towards lower temperatures while the opposite was observed for the corresponding mutant of AFC2 (H285Q). By contrast, the corresponding mutation in AFC1 (H304Q) had little effect on the temperature-activity profile. Taken together, these data start to unravel structural differences that allow distinct temperature-activity profiles of different AFCs and provide a framework for targeted mutations to alter the precise temperature response of the individual kinases.

A recent study suggested a negative regulatory role for AFC2 during thermomorphogenic growth in Arabidopsis^26^. According to that study, *afc2* single mutants displayed a slightly hyperresponsive hypocotyl elongation at 28 °C. In our hands, however, neither one of the two T-DNA insertion lines that had been used in that study (Supplementary Fig. 9c), nor our CRISPR/Cas9-generated *afc2* single mutant displayed hyperresponsive hypocotyl elongation. By contrast, our data indicate that *AFC1* and *AFC3* are required for warm temperature-dependent hypocotyl elongation since AFC2 could not compensate for the loss of AFC1 and AFC3 in the *afc1/3* double mutant. Consistently, warm temperature-dependent hypocotyl elongation could be partially restored by genetic complementation with *AFC3* in the triple mutant background thus clearly supporting a positive regulatory function of AFCs, in particular AFC1 and AFC3, during thermomorphogenesis. Beyond that, the triple mutant seedlings displayed WT-like elongation growth under skotomorphogenic conditions indicating that AFCs are specifically required for warm temperature-dependent growth responses. In addition, we did not observe major differences in the vegetative growth patterns between the WT and the two *afc1/2/3* triple mutant lines. However, we did note that both *afc1/2/3* triple mutant lines were late flowering (Supplementary Fig. 9d-f) suggesting a potential role for AFCs in the regulation of floral transition. Post-transcriptional regulation of flowering is well established for Arabidopsis and flowering time phenotypes have been reported for various mutants defective in splicing^38, 39, 40^. However, the functional role of AFCs in the regulation of floral transition was not examined any further in the course of this study.

Downstream of AFCs, our data support the notion that SR34 and SR34a are phosphorylation targets and furthermore suggest that these two SR proteins are involved in regulating hypocotyl elongation growth at elevated temperature. These hypotheses are based on three observations: First, immunodetection of phosphorylated SR proteins using an α-pan-phosphoepitope SR-specific antibody pinpointed towards AFC-dependent phosphorylation of SR34 and SR34a under our experimental conditions. Second, AFC-dependent phosphorylation of SR34a was confirmed *in planta* through immunoprecipitation-mass spectrometry. And third, *sr34* and *sr34a* mutant seedlings displayed decreased hypocotyl elongation growth at elevated temperature, resembling the *afc* triple knockout phenotype. Notably, the reduction in hypocotyl elongation growth in *sr34* and *sr34a* was less pronounced than in the *afc* triple mutants indicating an at least partially redundant function. However, further work is needed to unravel AFC target preferences and specificities in more detail, as well as precise targets of *sr34* and *sr34a* that then cause the hypocotyl elongation phenotype. Furthermore, the contribution of other SR proteins to warm temperature-dependent hypocotyl elongation cannot be ruled out.

This is the first report of a functional role of SR proteins in plant responses towards elevated temperatures. Thus far, SR34 and SR34a were only implicated in the context of ABA-dependent responses^41, 42^. However, SR proteins have been linked to various other abiotic stresses^21^, and they undergo extensive alternative splicing during heat shock themselves^16^. Based on these observations, a functional role in heat stress resistance has been hypothesised for SR proteins and AFC kinases^43^. The *afc1/2/3* triple mutant, however, did not show any differences in short- or long-term thermotolerance compared to the WT when exposed to heat stress above 37 °C, which is in agreement with the low *in vitro* kinase activities we observed for these temperatures.

For human SR proteins, a complex network of post-transcriptional auto- and cross-regulation has been shown^34, 35, 36^. Accordingly, SR proteins contribute to the post-transcriptional regulation of other SR protein transcripts^34^, but also regulate the splicing of their own pre-mRNA^35, 36^. In plants, several SR proteins were shown to regulate the splicing of other SR protein transcripts^41, 44, 45^ while post-transcriptional autoregulation has only been demonstrated for SCL33 and RS2Z33^46, 47^. In line with this, several SR protein transcripts were differentially spliced in the *afc* triple mutants pointing to a model in which AFCs regulate thermomorphogenic growth responses through first phosphorylating specific SR proteins. These in consequence alter alternative splicing and expression of additional members of the SR protein family, which then leads to the establishment of temperature-dependent alternative splicing patterns, gene expression and finally the thermomorphogenic growth responses (Fig. 5e). Prospective work needs to address the interaction between individual AFCs and SR proteins in more detail and determine their precise role in the regulation of warm temperature-dependent alternative splicing and target genes whose regulation controls thermomorphogenesis. This will be especially interesting, as our analysis of single and higher order *afc* mutants suggests a certain redundancy in controlling thermomorphogenesis, despite our finding that AFC activities are differentially affected at 28 °C.

Thus far, all known thermosensory mechanisms converge on PIF4/7-dependent signalling^48^. While we did not observe any enrichment of PIF4- or auxin-regulated genes among the differentially spliced genes in *afc* triple mutants in our RNA-Seq analysis, the induction of several *PIF4*-dependent marker genes was strongly reduced in *afc1/2/3* triple mutant seedlings. In addition, treatment of mutant seedlings with the synthetic auxin picloram resulted in a complete restoration of WT-like hypocotyl elongation growth at 28 °C. Together with the observation that the *pif4 afc1/2/3* quadruple mutant behaved identically to the *pif4* single mutant in terms of warm temperature-induced hypocotyl elongation, our data support a model by which AFCs act in the same signalling pathway as PIF4, likely upstream of auxin biosynthesis. The exact mechanism by which AFC-dependent alternative splicing interferes with PIF4-dependent signalling remains to be elucidated. In conclusion, this study provides compelling evidence that temperature-controlled AFC activity is evolutionarily conserved between plants and animals pointing towards a function of AFC kinases as molecular temperature sensors. Future studies will be needed, however, to clarify whether AFCs indeed conform to the proposed definition for plant thermosensors^49^. Nonetheless, our detailed understanding of the molecular mechanism underlying AFC temperature sensitivity provides a starting point for the targeted engineering of AFC isoforms with altered temperature-responsive properties. Given that AFCs are conserved across all land plants^24^, this may ultimately aid in the generation of crops with enhanced thermal resistance and the generation of temperature-switchable kinases might become a valuable tool for thermogenetics^50^.

## Methods

### Plant material and growth conditions

Mutant lines of *Arabidopsis thaliana* that were used in this study are listed in Supplementary Table 1. All mutants were in the Columbia-0 (Col-0) background unless stated otherwise. Seeds were surface sterilised using chlorine gas and sown on 0.5x Murashige and Skoog (MS) medium (pH 5.7, 1% (w/v) sucrose, 1.2% (w/v) phytoagar). Arabidopsis seeds were stratified for 24 to 48 hours at 4 °C in the dark and then transferred to a growth chamber at long-day conditions (16 h light/8 h darkness) set to 17 °C at 100-120 µmol photons m^-2^ s^-1^ (LED lights, Polyklima PK-520). For experiments with soil-grown plants, surface sterilised seeds were directly sown on soil, stratified for 48 hours at 4 °C and grown under the same conditions as described above.

### Three-dimensional protein structure analysis

To assess structural differences in the AFC family, predictions of full-length *A. thaliana* AFC1, AFC2 and AFC3 structures by AlphaFold^51^ published in the AlphaFold protein structure database^52^ as entries P51566 (AFC1), P51567 (AFC2) and P51568 (AFC3) were used. For analysis only the kinase domains, AFC1 residues 102-467, AFC2 residues 85-427 and AFC3 residues 58-400, were considered. Structure data was analysed with PyMOL 2.3.4 and Coot 0.9.6^53^. Similarity of structures was calculated with the rigid jFATCAT algorithm^54, 55^ via the RCSB PDB pairwise structure alignment tool.

### *In vitro* kinase assays

On-bead *in vitro* kinase assays were carried out as described previously^23^. 0.5 µM of the purified recombinant AFCs were pre-incubated with 2 µM GST-RS in reaction buffer (50 mM Tris-HCl pH 7.6, 10 mM MgCl_2_, 5 mM DTT, 0.1 mM spermidine) for 20 min at the indicated temperatures. After addition of an ATP mixture (γ-^32^P-ATP:ATP, 1:20k, ca. 0.3 Ci/mmol) to a final concentration of 22 µM the samples were incubated for 5 min at the same temperatures. The reaction was stopped by addition of 6x SDS sample buffer and incubation at 95°C for 5 min. Samples were analysed on 12% SDS-PAGE gels. Gels were stained with Coomassie, de-stained with 10% (v/v) acetic acid, 40% (v/v) ethanol and imaged. Subsequently gels were placed on filter paper, dried in a gel drier for 45 min and exposed on phosphoscreens overnight. The phosphoscreens were detected in a Typhoon FLA 7000 phosphoimager (650 nm laser, latitude L4, PMT = 500, 100 µm pixel size) and bands quantified using ImageQuant TL software. Details on the heterologous expression and purification of AFCs can be found in Supplementary Methods.

### Generation of CRISPR/Cas9-induced mutant lines

Genome-edited *afc* and *pif4* mutant lines were generated using CRISPR/Cas9^56^ as described previously^57^. Homozygous mutants exhibiting frame shifts and premature stop codons were identified by PCR amplification and sequencing of the targeted genomic regions. All primers used for the generation and sequencing of *afc* and *pif4* mutant lines are listed in Supplementary Table 2. More detailed information can be found in Supplementary Methods.

### Complementation of *afc1/2/3*

For complementation of the *afc1/2/3 #2* triple mutant, DNA was amplified from *A. thaliana* genomic DNA (Supplementary Table 2) to generate a product harbouring 1792 nt upstream of the *AFC3* start codon, the full predicted *AFC3* transcript and 447 nt downstream of the *AFC3* stop codon. The PCR product was cloned into pDONR221 (Thermo Fisher Scientific) and recombined into the plant transformation vector pMDC107 *via* Gateway cloning^58^. The *afc1/2/3 #2* triple mutant was transformed via Agrobacterium-mediated transformation and transgenic plants selected on medium containing Hygromycin. Presence of the transgene was confirmed by PCR and T2 seedlings were used for experiments. Primers used for the amplification of the genomic *AFC3* sequence are listed in Supplementary Table 2.

### Hypocotyl length measurements

For hypocotyl elongation assays, Arabidopsis seedlings were grown at 17 °C for four days before they were subjected to the indicated temperature treatments for another three days. For TG003and picloram treatments, Arabidopsis seedlings were grown for three days on 0.5x MS plates at 17 °C and then transferred to 0.5x MS plates containing 50 µM TG003 (Sigma-Aldrich, Catalogue No.: T5575), 1 µM picloram (Sigma-Aldrich. Catalogue No.: P5575) or an equivalent volume of DMSO as solvent control and grown for one (TG003) or two (picloram) additional days at 17 °C to allow for uptake of the compound. Subsequently, half of the plates were subjected to a three-day treatment at the indicated temperatures, whereas the other half remained at 17 °C as control. For all hypocotyl length measurements, plates were scanned at the end of the experiment using a flatbed scanner and hypocotyl lengths were quantified using the software Agnes Roots Measurements, Version 1.2.

### Quantitative real-time PCR

Total RNA was extracted from 7-day-old seedlings using a phenol-based RNA isolation method (NucleoZol, Macherey-Nagel, Catalogue No.: 740404) according to the manufacturer’s instructions. Following treatment with DNase I, cDNA was synthesised from 1 µg of RNA using RevertAid H minus reverse transcriptase (Thermo Scientific, Catalogue No.: EP0451) and oligo dT primers. Transcript abundance of target genes was determined through quantitative real-time PCR (qRT-PCR) using SYBR green as fluorescent dye (Thermo Scientific Absolute QPCR Capillary Mix, Catalogue No.: AB1285B). Transcript abundance of *SAND* (At2g28390) was used for normalisation. Primers used for qRT-PCR are listed in Supplementary Table 2.

### RNA-sequencing

To assess warm temperature-dependent splicing patterns of Col-0, *afc1/2/3 #1* and *afc1/2/3 #2*, seedlings were grown at 17 °C and then exposed to 28 °C for 24 hours starting at the end of day 7 or exposed to 28 °C for 8 hours starting on day 8. Control seedlings remained at 17 °C. Samples were taken at Zeitgeber Time (ZT) 11, i.e., 11 hours after the onset of light. Twenty seedlings were pooled per replicate and total RNA was extracted from homogenised plant tissue using a phenol-based RNA isolation method (NucleoZol, Macherey-Nagel, Catalogue No.: 740404). PolyA+ libraries were prepared by and sequenced at BGI Genomics (Hong Kong, China). Raw RNA sequence data was obtained in the FASTQ file format. For gene expression analysis, reads of each sample were mapped to the Arabidopsis genome (Col-0 TAIR10 genome release) using SALMON (v1.8.0)^59^ and quantified using DEseq2 (v1.28.1)^60^. For differential splicing analysis, raw reads were aligned to the Arabidopsis reference genome (Col-0 TAIR10 genome release) using STAR (version 2.7.5b)^61^. Quantification was done using rMATS (4.1.2)^62^. Pairwise comparisons were performed to identify differentially alternatively spliced transcripts between different genotypes or temperature conditions. Raw data from RNA-seq of this article can be found in Gene Expression Omnibus (GSE269859).

### Immunodetection of phosphorylated SR proteins

To assess warm temperature-dependent phosphorylation of SR proteins in Col-0 and *afc1/2/3 #2*, seedlings were grown at 17 °C for 9 days and then exposed to 28 °C for 1 hour. Control seedlings remained at 17 °C. Forty seedlings were pooled per replicate. Total protein was extracted from homogenised tissue, separated by SDS-PAGE and blotted onto PVDF membranes for immunodetection. Phosphorylated SR proteins were detected using a mouse monoclonal α-pan-phosphoepitope SR-specific antibody (1H4, Merck, Catalogue-No.: MABE50) at 0.025 µg/mL. Detection of histone H3 using a rabbit polyclonal α-H3 antibody (Agrisera, Catalogue No: AS10 710) at a dilution of 1:5000 served as loading control and was used for normalisation. Primary antibodies were detected using HRP-linked goat α-mouse IgG (Thermo Scientific, Catalogue No.: G-21040) and goat α-rabbit IgG (Merck, Catalogue No.: AP307F) antibodies at dilutions of 1:5000. Signal detection and quantification was achieved using the Clarity Western ECL substrate kit (Bio-Rad, Catalogue No.: 1705061), a ChemiDoc MP imaging system (Bio-Rad) and Image Lab, Version 6.1 (Bio-Rad laboratories).

### Immunoprecipitation-mass spectrometry

Coding sequences of *SR34a*, *AFC1*, *AFC2* and *AFC3* were amplified from Col-0 cDNA, cloned into *pDONR201* and then recombined into *pB7FWG2* and *pK7RWG2*, respectively, to yield GFP- and RFP-tagged constructs under the transcriptional control of the 35S promoter. *SR34a-GFP* was co-infiltrated with *RFP* or *AFC1-RFP*, *AFC2-RFP* or *AFC3-RFP* in 5-week-old *Nicotiana benthamiana* leaves. Infiltrated leaves were harvested after 3 days in 4 biological replicates for extraction of total protein. Enrichment of GFP-tagged SR34a, on-bead digestion and sample preparation were performed as described previously^63^. Samples were analysed via LC-MS/MS on an Ultimate 3000 RSLC nano LC (Thermo Fisher Scientific) in-line connected to a Q Exactive mass spectrometer (Thermo Fisher Scientific). Obtained MS/MS spectra were searched against the combined protein database of SR34a-GFP, AFC1-RFP, AFC2-RFP and AFC3-RFP, and using the *N. benthamiana* proteome downloaded from SolGenomics database containing 57,140 protein entries by the MaxQuant software (version v2.1.4.0). For SR34a-immunoprecipitated protein analysis all proteins that were detected in at least 3 replicates out of 4 were retained as reproducibly quantified proteins for statistical analysis. Details on the transient co-expression and mass spectrometry-based analysis can be found in Supplementary Methods.

### Statistical analyses

Unless stated otherwise data were analysed using two-factorial analysis of variance (ANOVA) followed by a TukeyHSD *post-hoc* test for pairwise comparisons. Normal distribution of the residuals was tested using the ‘plotresid’ function of the R package ‘RVAideMemoire’, Version 0.9-80^64^. Equality of variances was tested using Levene’s test. In case data did not meet the required assumptions, analyses were done on log-transformed data. All analyses were done using RStudio, Version 1.4.1717 and R, Version 4.1.1.

## Acknowledgments

We thank Prof. Dr. Dr. Martin J. Mueller (University of Wuerzburg, Germany) for constant support, Dr. Frank Waller (University of Wuerzburg, Germany) for proof-reading of the manuscript and Dr. Jan Draken (University of Wuerzburg, Germany) for excellent technical assistance with immunodetection. Andrea Rodi, Paul Wulf and Carlotta Wehrkamp contributed as rotation students. Seeds of *pif4-2* were kindly provided by Prof. Dr. Marcel Quint (Martin Luther University Halle-Wittenberg, Germany). We thank FUB-IT for computing time. This work was supported by the Universitätsbund Würzburg (to D.M.) and the German Research Foundation (to F.H., project number 270986915).

## Conflict of Interest

The authors declare that they have no conflict of interest.

## Notes

### Competing Interest Statement

The authors have declared no competing interest.

### Summary of Updates

Figures 3, 4 and 5 as well as 'Discussion' revised, author list and 'Supporting Information' updated

