## Supplementary Information for "AFC kinases regulate warm temperature-responsive growth in *Arabidopsis*"

**This PDF file includes:**

Supplementary Methods
Supplementary Tables 1 and 2
Supplementary Figures 1 to 9
Supplementary Information References

### Supplementary Methods

#### Heterologous expression and purification of AFCs

Full-length (f.l.) WT *AFC1-3* were amplified from *A. thaliana* cDNA and cloned into pGex-6P-1 vector for recombinant expression with an N-terminal GST-tag. Kinase domains (KDs) were amplified by PCR from these constructs and re-cloned into pGex-6P-1 vectors. Hybrid and point mutants were generated using 2-step PCRs and conventional cloning. The AFC3 f.l. Q254A mutant was generated via quick-change PCR. Recombinant proteins were expressed in *E. coli* BL21(DE3) pRare cultured in LB medium in the presence of 0.2 mM IPTG and incubated overnight at 18 °C. Bacterial pellets were resuspended in lysis buffer (50 mM Tris pH 7.2, 500 mM NaCl, 10 mM $MgCl_2$, 1 mM DTT), lysed via sonification and the lysate cleared by centrifugation. GST-AFC3 f.l. WT and Q254A were isolated by gravity flow glutathione agarose affinity chromatography using 5 mL of Protino® Glutathione Agarose 4B beads (Macherey-Nagel, Catalogue No.:745500). For elution lysis buffer including 20 mM reduced glutathione was used. Subsequently recombinant AFC3 was further purified by size exclusion chromatography (SEC) using a HiLoad Superdex S200 16/600 pg column (Cytiva GE28-9893-35) on an Äkta pure system (Cytiva). SEC buffer comprised 50 mM Tris pH 7.2, 300 mM NaCl, 10 mM $MgCl_2$, 1 mM DTT. AFCs for on-bead *in vitro* kinase assays were purified and recombinant synthetic substrate GST-RS was expressed and purified as previously described[1]. Oligonucleotides used for cloning of AFC coding sequences are listed in Supplementary Table 2.

#### Generation of CRISPR/Cas9-induced mutant lines

Mutants for *AFC1, AFC2*, *AFC3* and *PIF4* were generated using CRISPR/Cas9. In detail, target-specific guide sequences, close to the transcription start site of *AFC1* (exon 1), *AFC2* (exon 4), *AFC3* (exon 2) and *PIF4* (exon 1) were selected using the free online tool ChopChop[2]. To generate *afc* triple mutant lines guide sequences of *AFC1-3* were introduced in a sequential CRISPR array (each cassette composed of an U6 promoter, guide sequence spacer, guide scaffold and an U6 terminator) by PCR using respective *AFC* guide sequence containing oligonucleotides (given in Supplementary Table 2) and the vectors pCBC-DT1T2 and pCBC-DT2T3 as templates[3]. The final PCR product, resembling the guide expression units was introduced in the binary plant expression vector pHEE401E by GoldenGate cloning[3]. After plasmid integrity had been verified by sequencing, agrobacteria (GV3101) were transformed and subsequently used for floral-dip transformation. Following selection on 0.5x MS plates containing hygromycin at 50 mg/mL, successfully transformed progeny could be identified and genotyped by amplification of the *AFC* and *PIF4* target sites from genomic DNA and subsequent sequencing using the primers given in Supplementary Table 2. Due to egg-cell specific *Cas9* expression, homozygous *afc1-3* and *pif4* mutants could already be obtained in T1.

**Complementation of *afc1/2/3 #2***

For complementation of the *afc1/2/3 #2* triple mutant, DNA was amplified from *A. thaliana* genomic DNA with primers AFC3-geno-F and AFC3-geno-R (see Supplementary Table 2) to generate a product harbouring 1792 nt upstream of the *AFC3* start codon, the full predicted *AFC3* transcript and 447 nt downstream of the *AFC3* stop codon. The PCR product was cloned into pDONR221 (Thermo Fisher Scientific) and recombined into the plant transformation vector pMDC107 *via* Gateway cloning[4]. The *afc1/2/3 #2* triple mutant was transformed via Agrobacterium-mediated transformation and transgenic plants selected on medium containing Hygromycin. Presence of the transgene was confirmed by PCR and T2 seedlings were used for experiments.

**Petiole length measurements**

For petiole elongation assays, plants were grown on soil at 17 °C for 11 days before they were subjected to 28 °C for another ten days. Control seedlings remained at 17 °C. On day 21, the third leaf of all plants was cut at the basis and fixed on white paper using transparent sticky tape. The sheet of paper containing all leaves to be analysed was scanned using a flatbed scanner and petiole lengths were measured using Agnes Roots Measurements, Version 1.2.

**Flowering time**

To assess flowering time, seedlings were sown in potting soil, stratified at 4 °C in darkness for two days and then grown under long day conditions (16 h light/8 h darkness) at constant 17 °C for ten days. On day 11, plants were transferred to 21 °C or 28 °C or remained at 17 °C until the onset of flowering. Upon the onset of flowering, which was defined as the time point when the first flower had opened, the respective date and rosette leaf number were noted.

**Short-term acquired and long-term thermotolerance**

To determine their short-term acquired thermotolerance 14-day-old seedlings of Col-0 and *afc1/2/3 #2* grown on 0.5x MS agar plates as described above (16 h light/8 h darkness, 22 °C/20 °C) were acclimated for 2 hours at 37 °C, kept at 22 °C for two hours and then exposed to a 45 °C heat treatment for 2.5 h. As a control for the lethality of the heat shock two agar plates containing 15 seedlings of each genotype were exposed to the 45 °C heat treatment without prior acclimation. For long-term thermotolerance 11-day-old seedlings of Col-0 and *afc1/2/3 #2* grown on 0.5x MS agar plates as described above (16 h light/8 h darkness, 22 °C/20 °C) were exposed to constant 37 °C for four and six days, respectively. After the heat treatments agar plates were transferred back to control conditions, i.e., 22 °C. Seedling survival was assessed after 14 days based on their capacity to grew new leaves.

**Extraction and Quantification of TG003**

To determine endogenous TG003 levels in whole Arabidopsis seedlings, the roots of harvested seedlings were briefly rinsed with methanol to remove any TG003 that was adsorbed to the outer root surface, dried carefully, and the whole seedlings were flash frozen in liquid nitrogen. For extraction, approximately 10-20 mg of frozen plant powder was suspended in 500 µL of methanol and a zirconium oxide bead added. Following agitation in a bead mill (Retsch MM 400) for 5 min at 21 Hz, the samples were centrifuged at 15 000 x *g* and 4 °C for 5 min and 400 µL of the supernatants transferred to fresh reaction tubes. Subsequently, the samples were evaporated to dryness using a centrifugal evaporator and the dry residues were suspended in 25 µL of methanol/$H_20$ (50/50, v/v). Following centrifugation at 15 000 x *g* and 4 °C for 3 min, 20 µL of the supernatants were transferred to glass vials. TG003 was analysed using a Waters Acquity UPLC™ system equipped with an Acquity BEH C18 column (2.1 x 50 mm, 1.7 µm particle size equipped with a VanGuard pre-column and inline particle filter) coupled to a Waters 2996 photodiode array (PDA) detector and a Waters Quattro Premier triple quadrupole mass spectrometer. The following gradient program was employed at a flow rate of 0.25 mL/min: 5 to 100% B in 5 min, holding at 100% B for 2 min, and re-equilibration at 5% B for 3 min with solvent A consisting of water (0.1% formic acid, v/v) and solvent B consisting of acetonitrile. The injection volume was 5 µL and the column was operated at 40 °C. The identity of TG003 was confirmed based on $MS^2$-spectra generated by multiple reaction monitoring in positive electrospray ionisation mode using the following mass transitions: *m/z* 250 > 179, *m/z* 250 > 208, and *m/z* 250 > 222, respectively. Quantitation in plant extracts was achieved using the built-in PDA detector (λ = 362 ± 1.2 nm) of the UPLC™ system and based on an external calibration curve of pure TG003 injected at six different concentrations.

**Gene ontology term enrichment analysis**

Gene ontology (GO) term enrichment analysis of differentially alternatively spliced genes was done using the GO annotation dataset for 'Biological Process' within the PANTHER database (https://pantherdb.org/). Statistically significant overrepresentation of GO terms was tested employing Fisher's Exact test and FDR-based correction for multiple testing. Visualisation was done according to[5] following removal of redundant GO terms[6].

**Phosphoproteomics of Columbia-0 wild-type seedlings**

For phosphoproteomics of wild-type (Col-0) seedlings after exposure to elevated temperature, seedlings were grown on 0.5x MS-agar plates (pH 5.8, 1% (w/v) sucrose, 1% (w/v) agar) at 21 °C under continuous light conditions (100 µmol $m^{-2}$ $s^{-1}$ photosynthetically active radiation supplied by cool-white fluorescent tungsten tubes, Osram) for ten days. On day 11, plates were exposed to a temperature treatment at 28°C, followed by sample collection every 12 minutes until 60 minutes after high temperature treatment. Prior to the temperature treatment, a first set of samples ('0 min') was collected.

Total protein extraction and phosphopeptide enrichment was performed for four biological replicates as described previously[7]. The samples were analysed on an Ultimate™ 3000 RSLC nano LC (Thermo Fisher Scientific) connected to a Q Exactive orbitrap mass spectrometer (Thermo Fisher Scientific). Subsequently, MS/MS spectra were searched against the Arabidopsis database and MS1-based label-free quantification was performed with the MaxQuant software (version 1.5.4.1) from Orbitrap instruments[8, 9].

For quantitative phosphoproteomics, the 'Phospho(STY)sites' output files generated by the MaxQuant search were further analysed using 'PhosR' package in R[10]. For phosphoproteome data, only high-confidence hits with phosphorylation localisation probability > 0.75 were included in subsequent analyses. Statistics was down using Student's *t*-test and permutation-based false discovery rates (FDR) at $p<0.05$.

**Transient expression in tobacco and co-expression mass spectrometry**

*Agrobacterium tumefaciens* strain C58C1 containing the plasmids pB7FWG2::SR34a-GFP; pK7RWG2::AFC1-RFP, pK7RWG2::AFC2-RFP, pK7RWG2::AFC3-RFP was used for transient expression. *SR34a-GFP* was co-infiltrated with *RFP* or *AFC1-RFP*, *AFC2-RFP* or *AFC3-RFP* in 5-week-old *N. benthamiana* leaves. Infiltrated leaves were harvested after 3 days in four biological replicates. The finely ground material was suspended in homogenisation extraction buffer (50 mM Tris-HCl pH 8.0, 150 mM NaCl 0.5% and 0.5% NP40) containing appropriate amounts of the cOmplete™ protease inhibitor mixture (Roche, Catalogue-No.: 04693132001) and the PhosSTOP phosphatase inhibitor mixture (Roche, Catalogue-No.: PHOSS-RO). The samples were incubated by rotation at 4 °C for 30 min to maximise extraction of proteins. Cell debris was removed from the supernatant by centrifugation, and total protein from each sample was used for co-immunoprecipitation. Then 25 µL equilibrated GFP Trap (ChromoTek) magnetic beads per sample were added into total protein of each sample and incubated by rotation for 2 h at 4 °C. Beads were washed three times with wash buffer (50 mM Tris-HCl pH 7.5, 250 mM NaCl), followed by washing once with 1 mL 50 mM TEAB (pH 8.0). On-bead digestion was performed on the bound proteins with 0.5 µg trypsin (Promega) in 50 µL 50 mM TEAB (pH 8.0) for 2 h at 37 °C. The supernatants were retained, beads were washed twice, and the wash fractions were pooled with the supernatants. Samples were treated with 10 mM TCEP and 15 mM iodoacetamide (Thermo Fisher Scientific) for 30 min at 30 °C to reduce and carbamidomethylate cysteine residues. Then 0.5 µg trypsin were added to the samples for further digestion, and these were incubated overnight at 37 °C. The digestion was stopped by adjusting the sample to 1% TFA and samples were desalted using C18 Bond Elut tips (Agilent Technologies).

Samples were analysed via LC-MS/MS on an Ultimate 3000 RSLC nano LC (Thermo Fisher Scientific) in-line connected to a Q Exactive mass spectrometer (Thermo Fisher Scientific). The sample mixture was loaded on a trapping column (made in-house, 100 µm internal diameter (i.d.) × 20 mm, 5 µm C18 Reprosil-HD beads, Dr. Maisch, Ammerbuch-Entringen, Germany). After flushing from the trapping column, the sample was loaded on an analytical column (made in-house, 75 µm i.d. × 150 mm, 3 µm C18 Reprosil-HD beads, Dr. Maisch). Peptides were loaded with loading solvent A (0.1% TFA in water) and separated with a linear gradient from 98% solvent A' (0.1% formic acid in water) to 55% solvent B' (0.1% formic acid in water/acetonitrile, 20/80 (v/v)) over 170 min at a flow rate of 300 nL $min^{-1}$. This was followed by a 5 min wash reaching 99% of solvent B. The mass spectrometer was operated in data-dependent, positive ionisation mode, automatically switching between MS and MS/MS acquisition for the 10 most abundant peaks in a given MS spectrum. The source voltage was 3.4 kV and the capillary temperature was set to 275 °C. One MS1 scan (m/z 400–2000, AGC target 3 × 106 ions, maximum ion injection time 80 ms) acquired at a resolution of 70,000 (at 200 m/z) was followed by up to 10 tandem MS scans (resolution 17,500 at 200 m/z) of the most intense ions fulfilling predefined selection criteria (AGC target 5 × 104 ions, maximum ion injection time 60 ms, isolation window 2 Da, fixed first mass 140 m/z, spectrum data type: centroid, under fill ratio 2%, intensity threshold 1.7 × 104, exclusion of unassigned, 1, 5–8, > 8 charged precursors, peptide match preferred, exclude isotopes on, dynamic exclusion time 20 s). The HCD collision energy was set to 25% normalised collision energy and the polydimethylcyclosiloxane background ion at 445.120025 Da was used for internal calibration (lock mass).

Obtained MS/MS spectra were searched against the combined protein database of SR34a-GFP, AFC1-RFP, AFC2-RFP and AFC3-RFP, and using the *N. benthamiana* proteome downloaded from SolGenomics database containing 57,140 protein entries by the MaxQuant software (version v2.1.4.0) using the Ghent University High Performance Computing server. For SR34a-immunoprecipitated proteins analysis all proteins that were detected in at least three replicates out of four were retained as reproducibly quantified proteins for statistical analysis.

**Supplementary Table 1.** *Arabidopsis thaliana* genotypes used in this study.

| **Genotype** | **Description** | **Source** |
|---|---|---|
| Columbia-0 (Col-0) | Wild type | |
| *afc1* | *afc1* single mutant line | This study |
| *afc2* | *afc2* single mutant line | This study |
| *afc3* | *afc3* single mutant line | This study |
| *afc1/2* | *afc1 afc2* double mutant line | This study |
| *afc1/3* | *afc1 afc3* double mutant line | This study |
| *afc1/2/3 #1* | *afc1 afc2 afc3* triple mutant line | This study |
| *afc1/2/3 #2* | *afc1 afc2 afc3* triple mutant line | This study |
| *pif4 afc1/2/3 #2* | *pif4 afc1 afc2 afc3* triple mutant line | This study |
| *pif4* | *pif4* single mutant line | This study |
| *afc2-1* (Salk_118114) | T-DNA insertion line | NASC |
| *sr34* (Salk_102166) | T-DNA insertion line | NASC |
| *sr34a* (Salk_087841) | T-DNA insertion line | NASC |
| *pif4-2* (Sail_1288_E07) | T-DNA insertion line | Marcel Quint, Halle |

**Supplementary Table 2.** Oligonucleotides used in this study.

**CRISPR guide oligonucleotides**

| Gene | Locus | Primers (5'-3') |
|---|---|---|
| *AFC1* | AT3G53570 | ATTGCAGCATCCCAAGTCAATCG<br>AAACCGATTGACTTGGGATGCTG |
| *AFC2* | AT4G24740 | ATTGGGTGTGAAGAAATACCGTG<br>AAACCACGGTATTTCTTCACACC |
| *AFC3* | AT4G32660 | ATTGAGGACATGGAAGCGATGGG<br>AAACCCCATCGCTTCCATGTCCT |
| *PIF4* | AT2G43010 | ATTGGATCTGACATGGAACACCA<br>AAACTGGTGTTCCATGTCAGAGTC |

**Genotyping primers for CRISPR/Cas9-induced mutants**

| Gene | Locus | Primers (5'-3') |
|---|---|---|
| *AFC1* | AT3G53570 | AGGAATGTTGAATTCCCTCATC<br>CGGAGGTTGAAATACCTGAAAG |
| *AFC2* | AT4G24740 | ATAGCAAAATGGGGGAAGGTAT<br>ACAGTACTTACCGATTGCCACC |
| *AFC3* | AT4G32660 | CAGAGGTTGCTCCTCTTTCTTC<br>AGGAGTGAGATTGTCCCTCAAA |
| *PIF4* | AT2G43010 | TGCTCCTCCTTGTATCTCTTAATCA<br>TGTGGCCTAGACATCAATACACA |

**Primers used for cloning of *AFC3* genomic sequence for complementation**

| Gene | Locus | Primers (5'-3') |
|---|---|---|
| *AFC3* | AT4G32660 | GTCCCAAAAGCAGGCAAGAG<br>GGTCGATTTCACTGAAGATCTC |

**Genotyping primers for T-DNA insertion lines**

| Genotype | Locus | Primers (5'-3') |
|---|---|---|
| *afc2-1* | AT4G24740 | TCTTTAACCACGCTCCAAGTG<br>AGATCAAATTTGGTTCACCATC |
| *sr34* | AT1G02840 | CTAACAACAAGCCAGTTTCGG<br>GATCTTGCCAAGAAGCAGATG |
| *sr34a* | AT3G49430 | TAATGTCACCGGGCAAGTTAC<br>TTGTTGGCTTCAGACCAAATC |

**RT-qPCR primers**

| Gene | Locus | Primers (5'-3') |
|---|---|---|
| *SAND* | AT2G28390 | GGCCTTTTTGCATGCCTATGT<br>AAAGTCCAAAGGGACCTCCG |
| *PIF4* | AT2G43010 | ATCATCTCCGACCGGTTTGC<br>AGTGGCTCACCAACCTAGTG |
| *YUCCA8* | AT4G28720 | CGATGAGACCAGTGGCTTGT<br>CGTGAATCACCTCACCGGAA |
| *SAUR19* | AT5G18010 | CTTCAAGAGCTTCATAATAATTCAAACTT<br>GAAGGAAAAAATGTTGGATCATCTT |
| *SAUR20* | AT5G18020 | AACTTGAATCTTTTCATACATCTTCAGAAGA<br>TAACTAGGAAGAAAAATGTTGGCTCATC |
| *SR34* | AT1G02840 | CATCTAGGAGATCAGAGTTTCGTG<br>GTCGAGCTTTTTCAGCGCAT |
| *SR34a* | AT3G49430 | GTTGAGCTTGCACATGGTGG<br>GATGGGAGCCCACGTACAAT |

**Primers used for RT-PCR of *SR45a* transcript isoforms**

| Gene | Locus | Primers (5'-3') |
|---|---|---|
| *SR45a* | AT1G07350 | TGTCCTGGACCCATGGACTAG<br>GACTTATGTCGTCCTCGAGCAG |

**Primers used for full-length transcript detection**

| Gene | Locus | Primers (5'-3') |
|---|---|---|
| *SR34* | AT1G02840 | AATTGCGGAGGCTGAGAGAT<br>TCAGAAGGAGGGAAAAGAAACGA |

**Primers used for cloning of AFC coding sequences for *in vitro* assays**

| Gene | Locus | Primers (5'-3') |
|---|---|---|
| *AFC1-FL* | AT3G53570 | GGATCCATGCAAAGCAGTGTGTATC<br>CTCGAGTTAGTTCTTTTGGTTGTATAAAAATG |
| *AFC1-KD* | AT3G53570 | ATATGGATCCTATGTCTTTGTTGTTGGGGA<br>CTCGAGTTAGTTCTTTTGGTTGTATAAAAATG |
| *AFC2-FL* | AT4G24740 | ATATGGATCCGAGATTATGGAGATGGAGCG<br>ATATCTCGAGTCAAATTCAAGTCCCGCTA |
| *AFC2-KD* | AT4G24740 | ATATGGATCCCATTACATATTTGAACTGGGAG<br>ATATCTCGAGTCAAATTCAAGTCCCGCTA |
| *AFC3-FL* | AT4G32660 | ATATGGATCCATGATAGCTAACGGATTCGAG<br>ATATCTCGAGTCAACTTGAGCTCTTAAAGAAAG |
| *AFC3-KD* | AT4G32660 | ATATGGATCCGATCGTGATGGTCATTATGT<br>ATATCTCGAGTCAACTTGAGCTCTTAAAGAAAG |

FL: full length
KD: kinase domain

**Primers used for AFC mutant generation for *in vitro* assays**

| Gene | Primers (5'-3') |
|---|---|
| AFC1-H304Q | CTACATTGTATCCACAAGACAGTACCGCGCACCAGAAGTTATCCTAGG<br>CCTAGGATAACTTCTGGTGCGCGGTACTGTCTTGTGGATACAATGTAG |
| AFC2-H285Q | CCTACATTGTATCAACCAGACAGTATAGGGCACCAGAAGTCATTTTAGG<br>CCTAAAATGACTTCTGGTGCCCTATACTGTCTGGTTGATACAATGTAGG |
| AFC3-H257Q | CACTCCATTGTCCAAACAAGACAGTACAGATCCCCCGAGGT<br>ACCTCGGGGGATCTGTACTGTCTTGTTTGGACAATGGAGTG |
| AFC3-Q254A | GGATTCATCACTCCATTGTCGCAACAAGACATTACAGATCCC<br>GGGATCTGTAATGTCTTGTTGCGACAATGGAGTGATGAATCC |
| AFC3-KD-AFC2-AS_1 | GATTTTGGCAGTACTACTTATGAGCGCCAGGACCAAACCTACATTGTCTCAACAAGACATTACAG<br>CTGTAATGTCTTGTTGAGACAATGTAGGTTTGGTCCTGGCGCTCATAAGTAGTACTGCCAAAATC |
| AFC3-KD-AFC2-AS_2 | CAAGACATTACAGAGCACCCGAGGTCATTTTAGG<br>CCTAAAATGACCTCGGGTGCTCTGTAATGTCTTG |

**Primers used for cloning of coding sequences for co-expression in *Nicotiana benthamiana***

| Gene | Primers (5'-3') |
|---|---|
| *AFC1* | GGGGACAAGTTTGTACAAAAAAGCAGGCTTCATGCAAAGCAGTGTGTATC<br>GGGGACCACTTTGTACAAGAAAGCTGGGTCGTTCTTTTGGTTGTATAAAAATGG |
| *AFC2* | GGGGACAAGTTTGTACAAAAAAGCAGGCTTCATGGAGATGGAGCGTGTG<br>GGGGACCACTTTGTACAAGAAAGCTGGGTCTCTTCTCCTTGCGAAAAACG |
| *AFC3* | GGGGACAAGTTTGTACAAAAAAGCAGGCTTCATGATAGCTAACGGATTCGAGAG<br>GGGGACCACTTTGTACAAGAAAGCTGGGTCACTTGAGCTCTTAAAGAAAGGATGG |
| *SR34a* | GGGGACAAGTTTGTACAAAAAAGCAGGCTTCATGAGTGGGCGATTTTCTCGG<br>GGGGACCACTTTGTACAAGAAAGCTGGGTCCACACTGCCTTCGCGAACC |

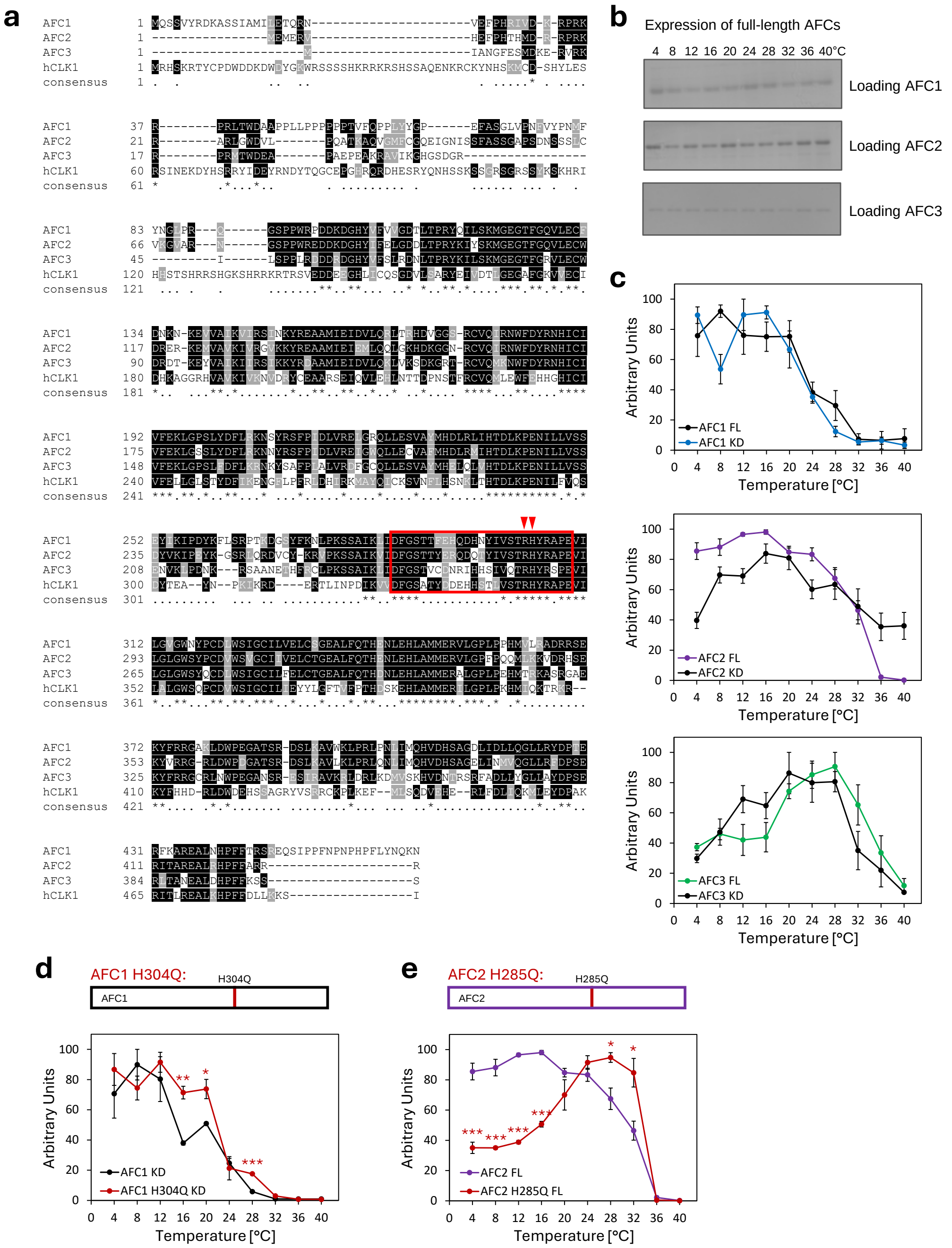


**Supplementary Figure 1.** **(a)** Alignment of the amino acid sequences of Arabidopsis AFC1, AFC2 and AFC3, and human CLK1. The consensus sequence is indicated below. Asterisks denote conservation of individual amino acids across all four sequences. The activation segment is indicated by a red box. Red arrows indicate the conserved arginine and histidine residues that are instrumental for the temperature sensitivity of human CLK1. **(b)** AFC kinase loading controls for *in vitro* assays shown in Fig. 1b as determined by Coomassie staining. **(c)** Comparison of temperature-dependent kinase activities between recombinant full-length (FL) AFCs and the respective kinase domains (KD) only. For quantification, the highest signal intensity within an assay was set to 100 and used for normalisation. Shown are mean values ± SE; $n$ = 3 and $n$ = 4 for AFC1 FL and KD, respectively, $n$ = 6 and $n$ = 10 for AFC2 FL and KD, respectively, and $n$ = 3 for AFC3 FL and KD, respectively. **(d,e)** Temperature-dependent activity of AFC1 H304Q (d) and AFC2 H285Q (e) point mutants. Shown are mean values ± SE from $n$ = 3 and $n$ = 5 replicates for AFC1 KD and H304Q KD, respectively, and $n$ = 3 and $n$ = 6 replicates for AFC2 FL and H285Q FL, respectively. Statistically significant differences were determined by Student's $t$-tests (*: $p < 0.05$, **: $p < 0.01$, ***: $p < 0.001$). Note that data for AFC2 FL depicted in (e) are the same as presented in (c). FL: full-length, KD: kinase domain.

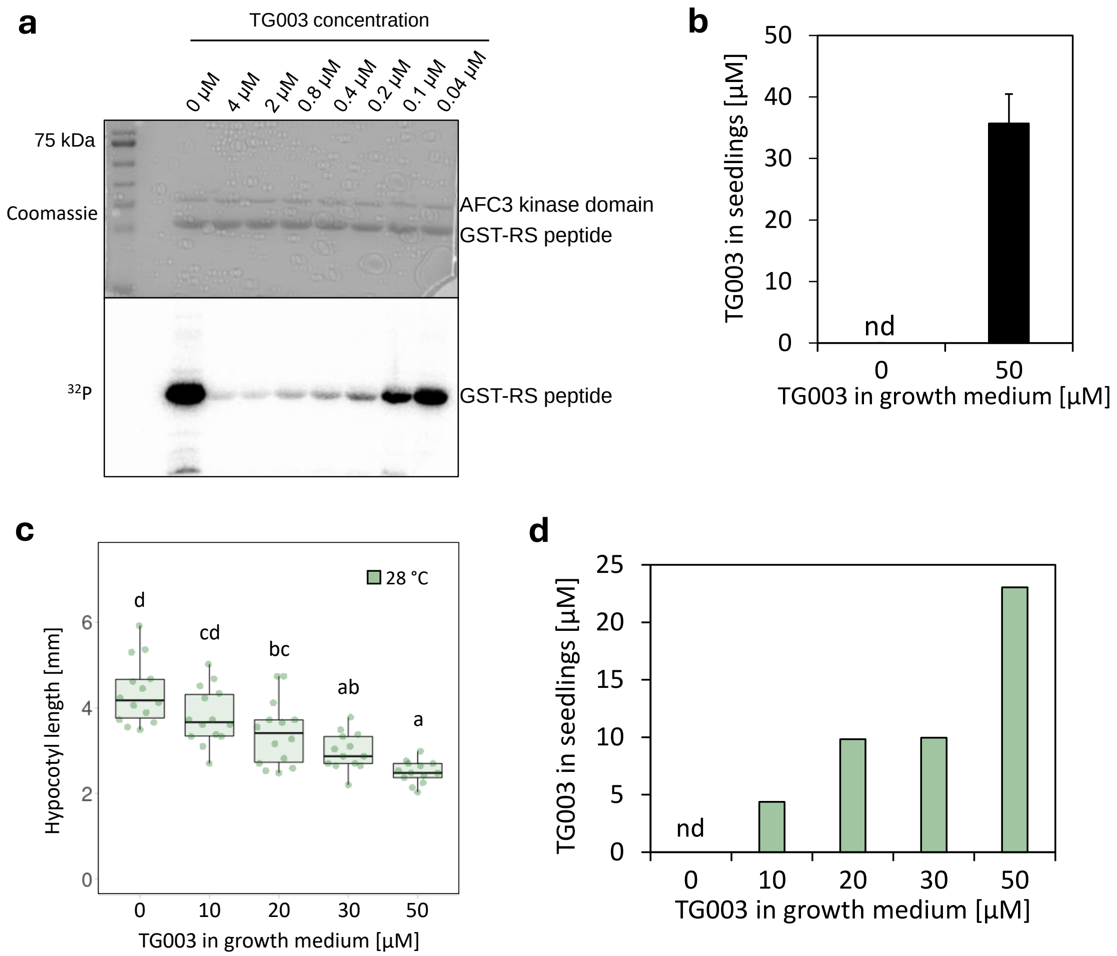


**Supplementary Figure 2. (a)** *In vitro* activity of recombinant AFC3 in the presence of different concentrations of the inhibitor TG003. Shown are the Coomassie-stained gel (top) and the radiograph showing phosphorylation of the SR peptide (bottom, $P^{32}$). **(b)** Determined TG003 concentration in Col-0 seedlings grown on 0.5x MS-Agar medium supplemented with 50 µM TG003 or without TG003 that were used for hypocotyl length measurements shown in Fig. 3A ($n$ = 4-6). TG003 was detected by UPLC-PDA; nd: not detected. **(c)** Hypocotyl lengths of seven-day-old Col-0 seedlings that were grown in the presence of different TG003 concentrations and exposed to 28 °C for three days ($n$ = 13-14 seedlings per treatment) **(d)** Determined TG003 concentrations in Col-0 seedlings grown on 0.5x MS-Agar supplemented with different concentrations of TG003 from (c) ($n$ = 1, note that for each concentration all seedlings had to be pooled to facilitate TG003 detection). TG003 was detected by UPLC-PDA; nd: not detected. Statistically significant differences were determined by one-way ANOVA followed by a Tukey HSD *post-hoc* test and are indicated by different letters above boxes ($p$ < 0.05).

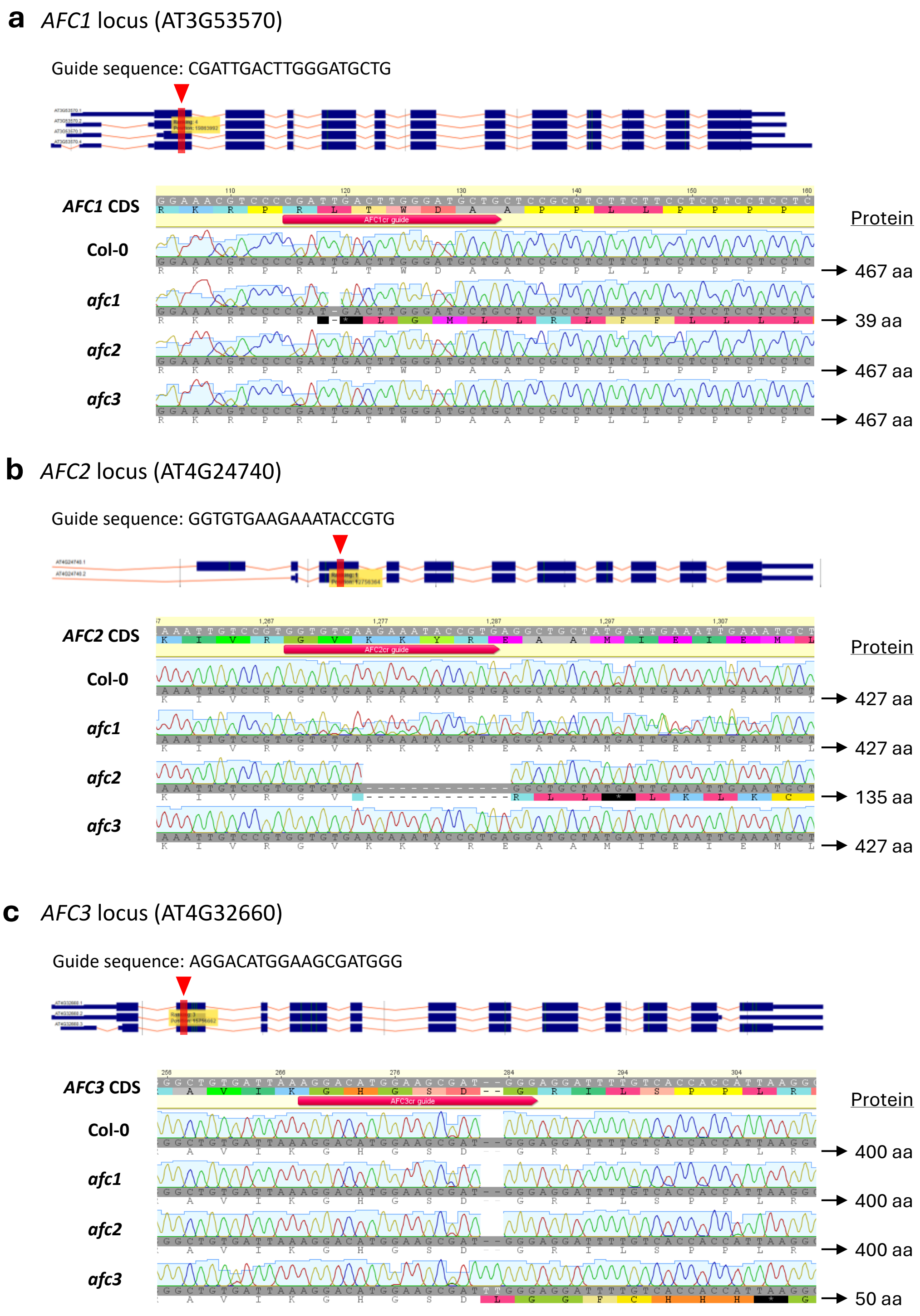


**Supplementary Figure 3. Characterisation of the CRISPR/Cas9-induced mutation alleles in the generated *afc* single mutant lines. (a-c),** Schematic representations of the coding regions for the different known isoforms of *AFC1* (a), *AFC2* (b) and *AFC3* (c) are shown above the respective sequencing results for Col-0 as well as *afc1*, *afc2* and *afc3* single mutant lines. Exons are given in blue and introns as red lines. Suited CRISPR guide sequences were identified by ChopChop. The position of the used CRISPR guide sequences are indicated by red triangles and boxes. Sequencing revealed deletion of a single thymine in *AFC1* in the *afc1* single mutant line (a), a 13 bp-deletion in *AFC2* in the *afc2* mutant line (b) and insertion of two thymines in *AFC3* in the *afc3* single mutant line (c). WT-like alleles for the non-addressed loci were confirmed for all three single mutant lines. In all cases, the modifications lead to frame shifts resulting in the occurrence of premature translation termination codons (the resulting protein lengths are indicated next to the chromatograms). aa: amino acid

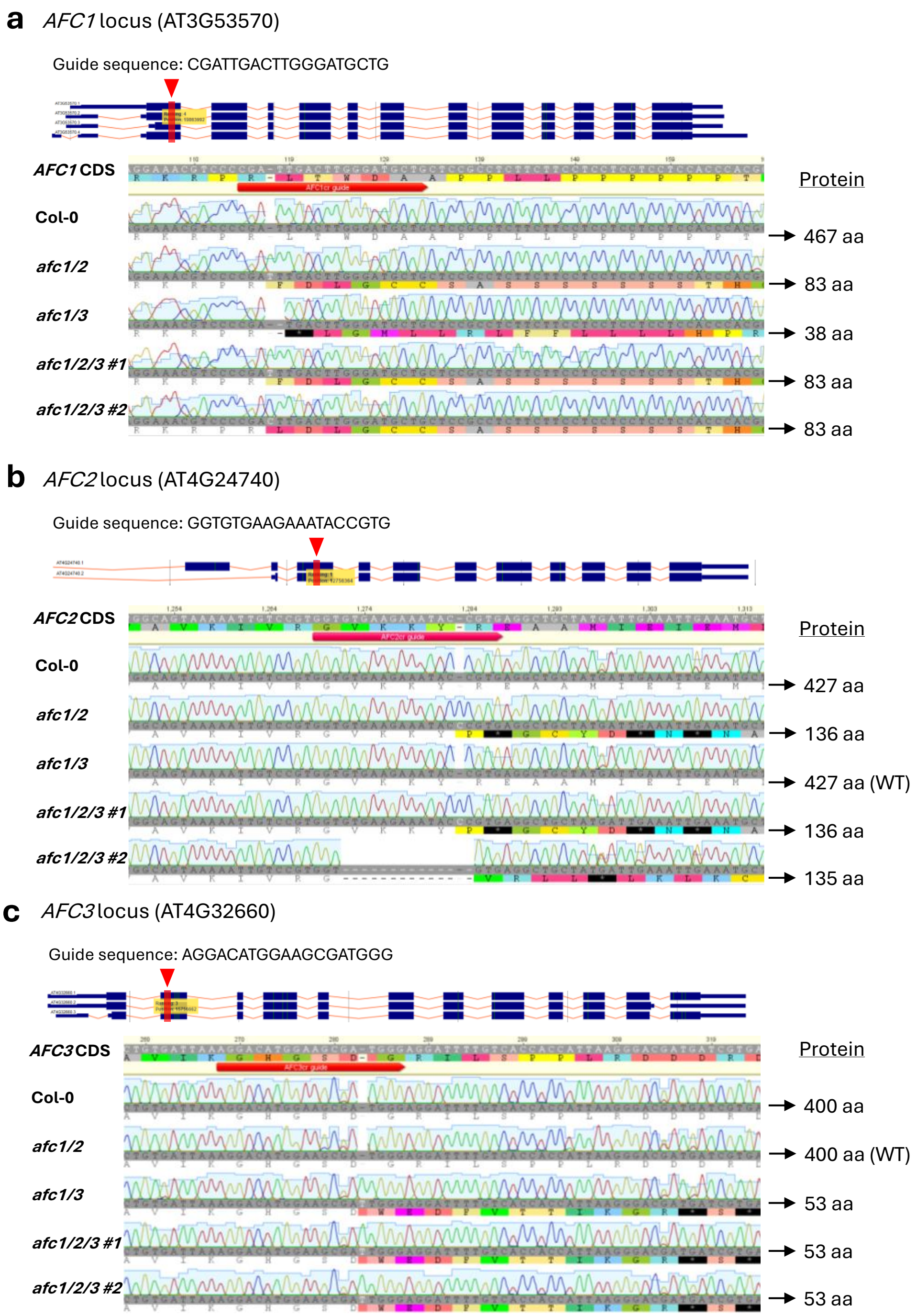


**Supplementary Figure 4. Characterisation of the CRISPR/Cas9-induced mutation alleles in the generated *afc* double and triple mutants. (a-c)** Schematic representations of the coding regions for the different known isoforms of *AFC1* (a), *AFC2* (b) and *AFC3* (c) are shown above the respective sequencing results for Col-0, the *afc1 afc2* (*afc1/2*) and *afc1 afc3* (*afc1/3*) double, and the two *afc1 afc2 afc3* triple mutant lines (*afc1/2/3 #1* and *afc1/2/3 #2*). Exons are given in blue and introns as red lines. Suited CRISPR guide sequences were identified by ChopChop. The position of the used CRISPR guide sequences are indicated by red triangles and boxes. For *AFC1*, sequencing revealed the insertion of a single thymine or cytosine in *afc1/2*, *afc1/2/3 #1* and *afc1/2/3 #2*, respectively, while for *afc1/3* the deletion of a single thymine was detected (a). For *AFC2*, insertion of a single cytosine and a 13 bp-deletion were detected for *afc1/2*, afc1/2/3 #1 and *afc1/2/3 #2*, respectively, while for *afc1/3* the wild-type allele was detected (b). For *AFC3*, insertion of a single thymine was detected for *afc1/3*, afc1/2/3 #1 and *afc1/2/3 #2*, respectively, while for *afc1/2* the wild-type allele was detected (c). In all cases, the modifications lead to frame shifts resulting in the occurrence of premature translation termination codons (the resulting protein lengths are indicated next to the chromatograms). aa: amino acid

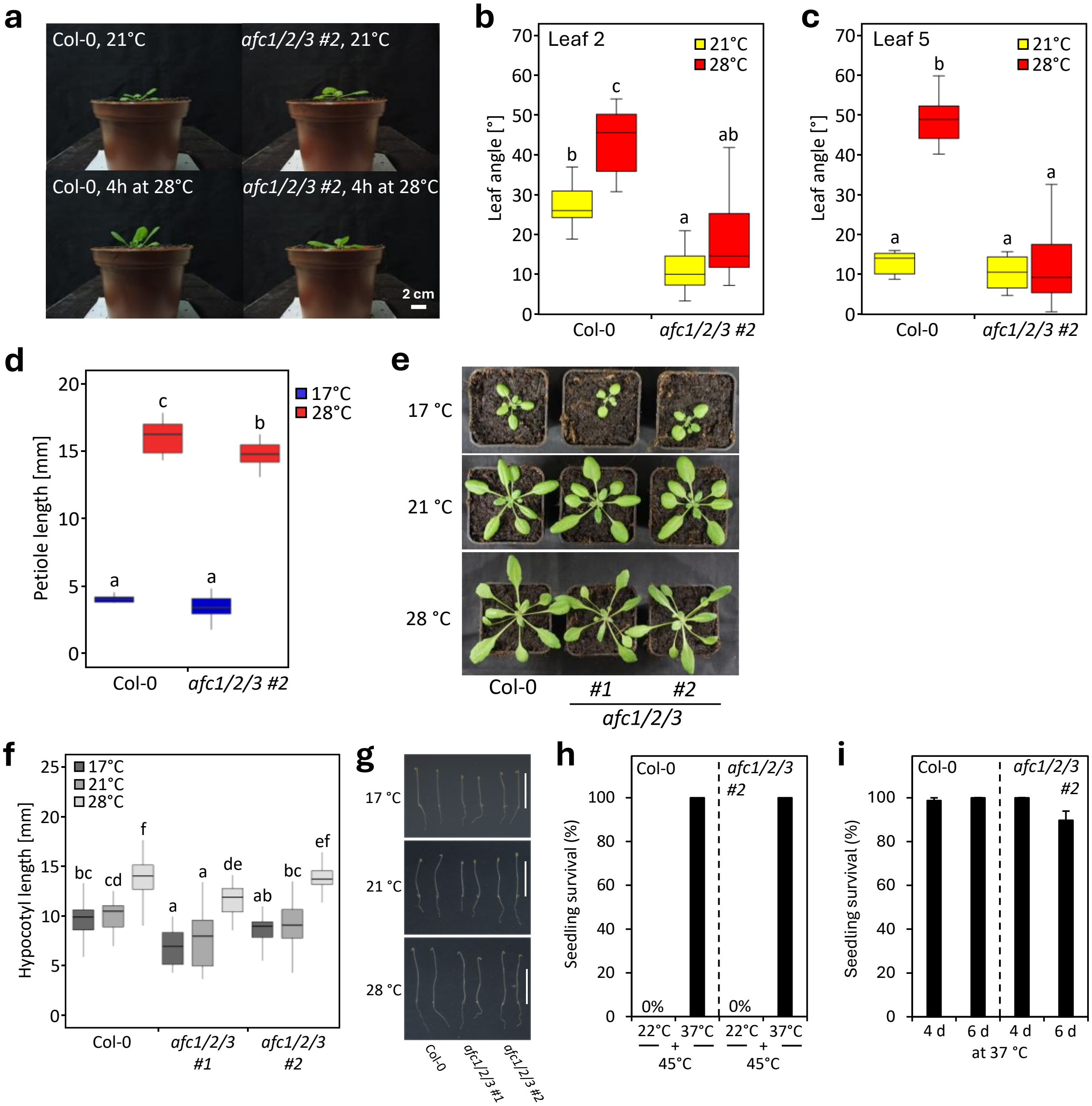


**Supplementary Figure 5. Phenotypic characterisation of *afc1/2/3* triple mutant plants. (a)** Representative pictures of Col-0 and *afc1/2/3 #2* plants that were used for leaf angle measurements shown in (b) and (c). Scale bar: 2 cm. **(b,c)** Leaf angles of leaf two (c) and leaf five (d) of four-week-old Col-0 and *afc1/2/3 #2* plants that were exposed to 28 °C for four hours or kept at control conditions, i.e., 21 °C (*n* = 7-8 plants per genotype and condition). **(d)** Petiole lengths of the third leaves of three-week-old Col-0 and *afc1/2/3 #2* plants at 17 °C and 28 °C. Plants were grown at 17 °C for 11 days and then exposed to the indicated temperatures for another ten days (*n* = 9-10 plants per genotype and treatment). **(e)** Representative pictures of 23-day-old Col-0, *afc1/2/3 #1* and *afc1/2/3 #2* plants grown under long-day conditions at different temperatures. Plants were grown for ten days at 17 °C and then for another 13 days at the indicated temperatures. Scale bar: 1 cm. **(f,g)** Hypocotyl lengths (f) and representative pictures (g) of dark-grown seedlings of Col-0, *afc1/2/3 #1* and *afc1/2/3 #2* that were grown at 17 °C, 21 °C or 28 °C. For 17 °C, hypocotyl lengths were measured after 6 days. For 21 °C and 28 °C, hypocotyl lengths were measured after 4 days (*n* = 17-41 seedlings per genotype and condition). **(h,i)** Short-term acquired (I) and long-term (i) thermotolerance of Col-0 and *afc1/2/3 #2* seedlings (mean ± SE, *n* = 5-7). See Supplementary Methods for more detailed information. Statistically significant differences were determined by two-way ANOVA followed by a Tukey HSD *post-hoc* test and are indicated by different letters above boxes (*p* < 0.05).

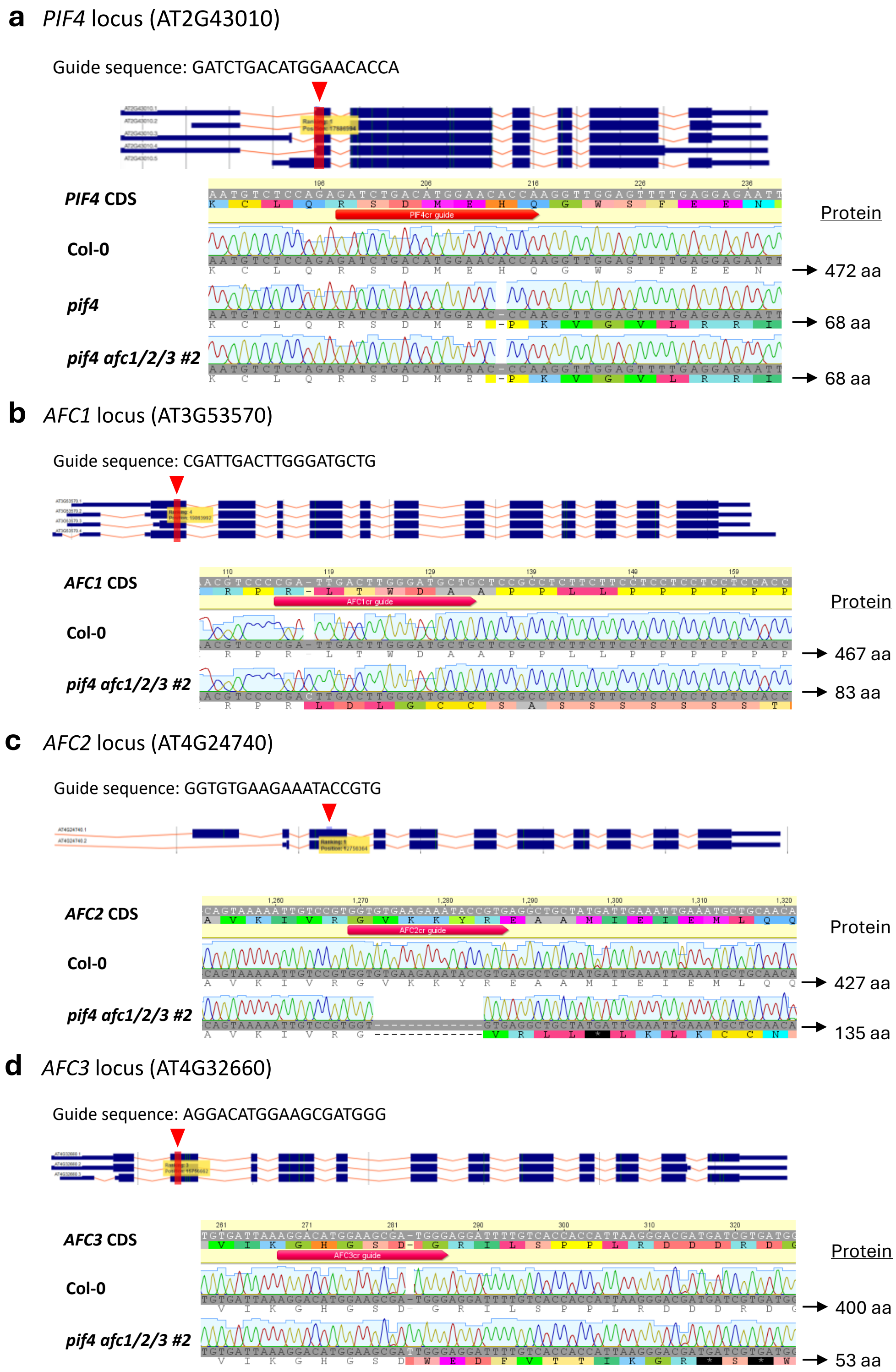


**Supplementary Figure 6. Characterisation of the CRISPR/Cas9-induced mutation alleles in the generated *pif4* mutant lines. (a)** Schematic representation of the coding regions for the different known isoforms of *PIF4*. Sequencing of the *PIF4* locus revealed deletion of a single adenine in *pif4* (Col-0 background) and the *pif4 afc1/2/3 #2* quadruple mutant. Exons are given in blue and introns as red lines. Suited CRISPR guide sequences were identified by ChopChop. The position of the used CRISPR guide sequences are indicated by red triangles and boxes. **(b-d)** Sequencing of the *AFC1* (b), *AFC2* (c) and *AFC3* (d) loci confirmed the expected mutations (see Supplementary Fig. 4) present in *afc1/2/3 #2* used for the generation of the quadruple mutant. In all cases, the modifications lead to frame shifts resulting in the occurrence of premature translation termination codons (the resulting protein lengths are indicated next to the chromatograms). aa: amino acid

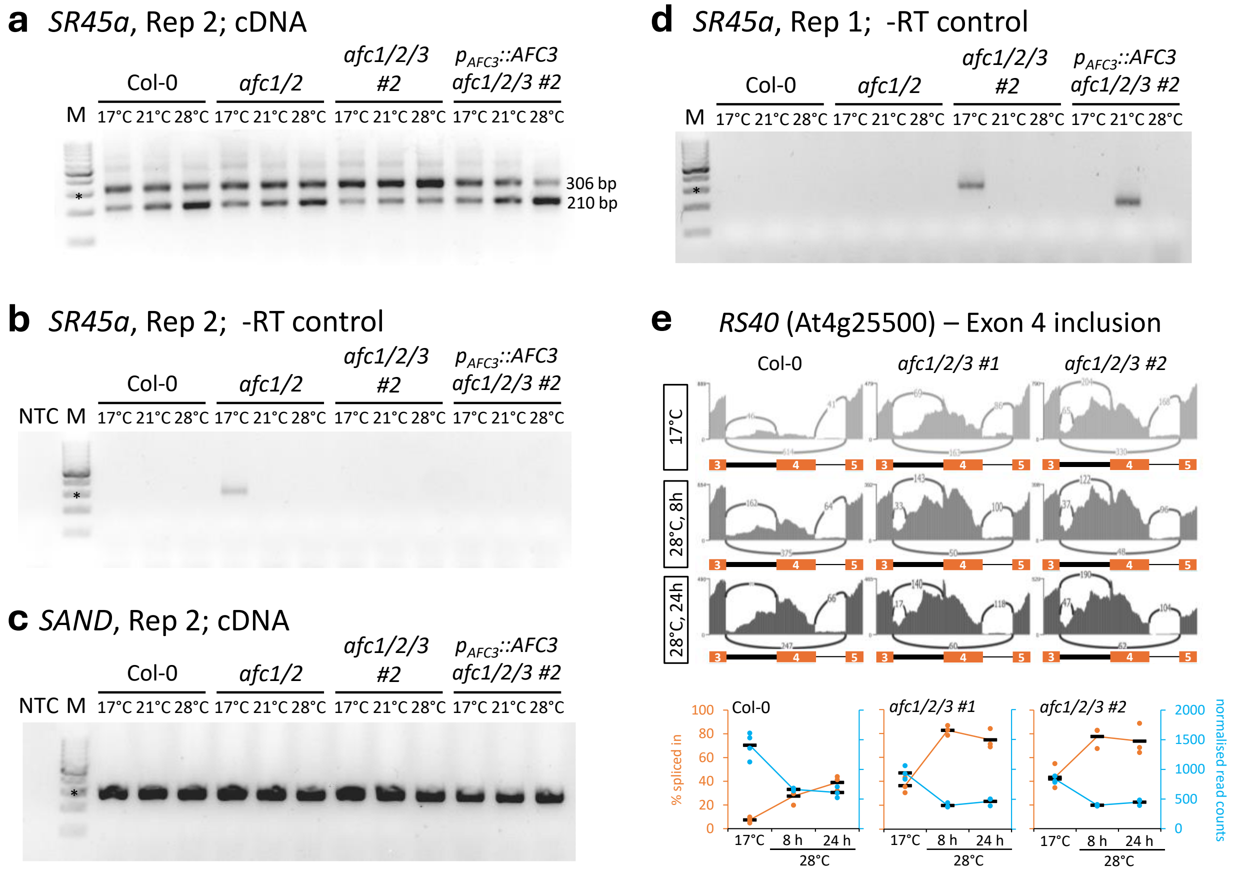


**Supplementary Figure 7. (a)** Detection of two distinct *SR45a* splicing isoforms in Col-0, *afc1/2*, *afc1/2/3 #2* and *afc1/2/3 #2* complemented with genomic *AFC3* ($p_{AFC3}$::*AFC3 afc1/2/3 #2*) by reverse transcription PCR. **(b)** Minus reverse transcription control using the same *SR45a*-specific primer pair as in (a) and the same RNA samples without prior cDNA synthesis. **(c)** Expression control using the same cDNA samples as in (A) and a *SAND* (At2g28390)-specific primer pair. **(d)** Minus reverse transcription control related to Fig. 4G using the same RNA samples without prior cDNA synthesis and an *SR45a*-specific primer pair. Seedlings were grown at 17 °C for four days and then exposed to the indicated temperatures for another three days. We noted minor impurities in isolated -RT lanes. The products are much less intense than in the reactions containing the cDNA. These impurities therefore do not affect the interpretation of these data. Asterisks in the DNA marker lane indicate 300 bp length. M: DNA marker. NTC: non-template control **(e)** Temperature-dependent splicing of *RS40* exon 4 in Col-0, *afc1/2/3 #1* and *afc1/2/3 #2* as determined by RNA-Seq. Inclusion of exon 4 produces a poison isoform containing two premature termination codons. Representative Shashimi plots for the indicated temperatures are shown on top. rMATS-derived percentages of exon 4 inclusion (orange) and *RS40* normalised read counts (blue) are shown below (n = 3-4). Horizontal lines represent mean values.

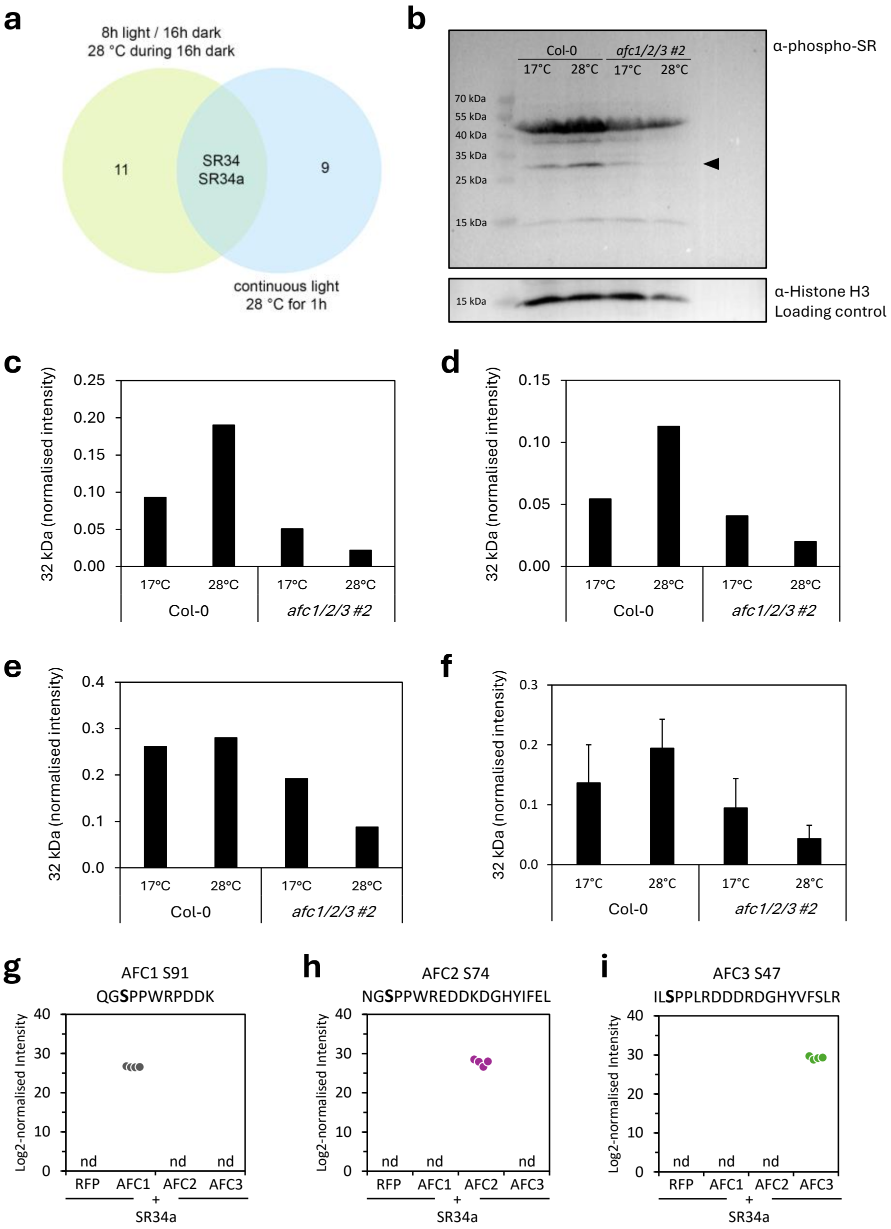


**Supplementary Figure 8. (a)** Venn diagram showing the intersection between warm temperature-regulated SR protein phosphosites in Col-0 seedlings exposed to 28 °C from two independent data sets (see Supplementary Data 5a,b). The experimental conditions are described in brief next to the Venn diagram and in detail in the Supplementary Methods above. **(b)** Representative immunoblot images of phospho-SR protein (top) and histone H3 (bottom) detection in identical samples. Proteins were isolated from nine-day-old seedlings of Col-0 and *afc1/2/3 #2* that were exposed to 28 °C for one hour or remained at 17 °C as control. SR protein phosphorylation was assayed using an α-pan-phosphoepitope SR-specific antibody and normalised to histone H3 signal intensity as detected by an α-H3 antibody. **(c-e)** Quantification of temperature-dependent SR protein phosphorylation of the protein band at 32 kDa (indicated by an arrow in (a)) from three individual immunoblot experiments. Signal intensities were normalised to the respective H3 signal intensity. **(f)** Mean values ± SE from experiments shown in (c-e), *n* = 5 biological replicates per genotype and temperature. **(g-i)** Log2 intensities of AFC1 (g), AFC2 (h) and AFC3 (i) phosphopeptides containing the indicated phosphorylated serine residues (bold) as determined by co-immunoprecipitation mass spectrometry following co-expression of SR34a-GFP with RFP-tagged AFC1, AFC, AFC3 or free RFP in *N. benthamiana* leaves. Immunoprecipitation of GFP-tagged SR34a was achieved using GFP trap magnetic beads. Horizontal lines indicate mean values (*n* = 4 replicates). nd: not detected

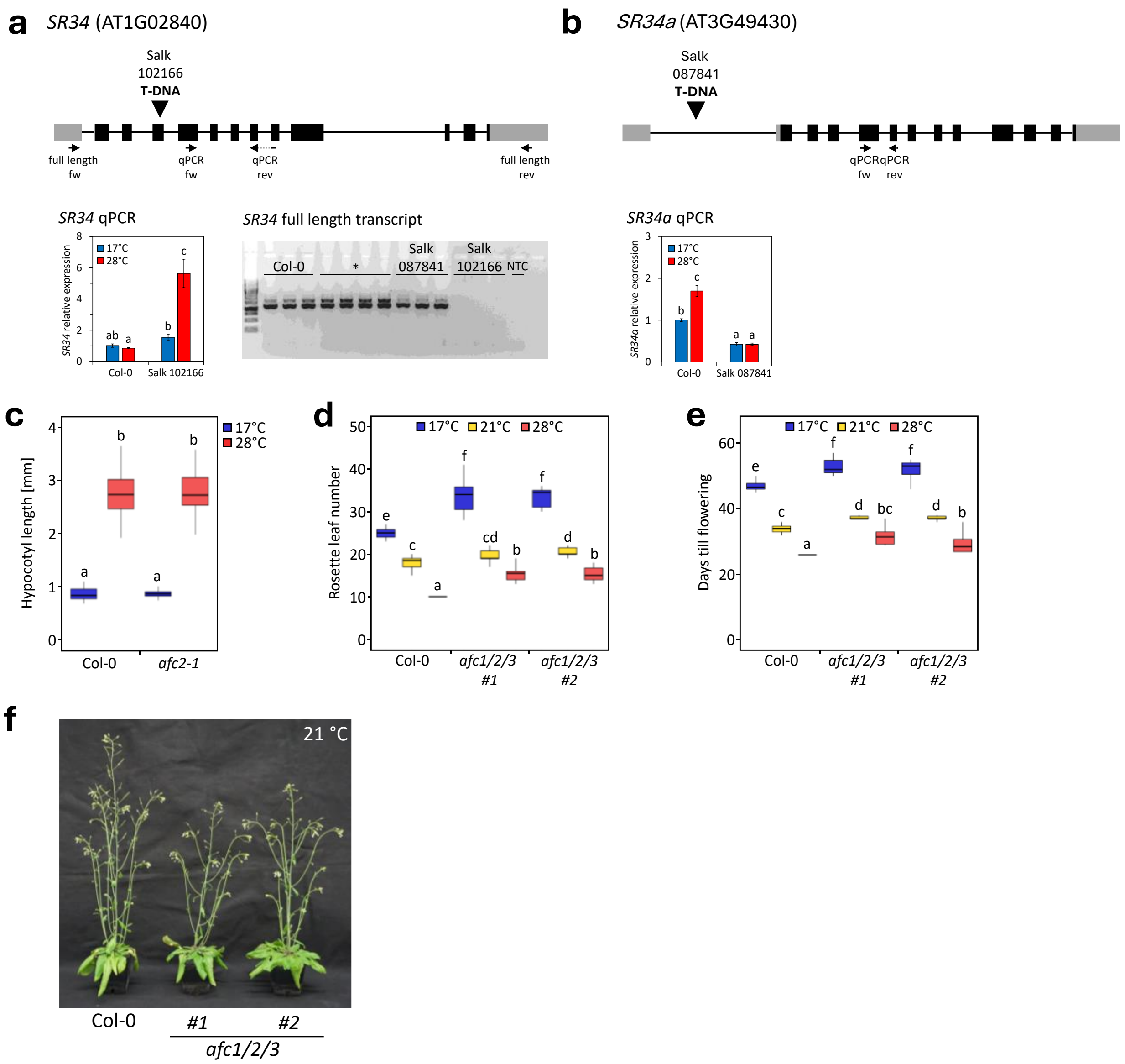


**Supplementary Figure 9. (a,b)** Characterisation of T-DNA insertion lines for *SR34* (a) and *SR34a* (b). Depicted are schematic representations of the two genomic regions with exons and introns indicated as boxes and lines, respectively. Grey boxes indicate untranslated regions. Insertion sites are indicated above, and primers used for RT-qPCR or amplification of full-length coding sequences from cDNA are shown below. Temperature-dependent transcript abundance is shown for seven-day-old Col-0 wild-type and the respective mutant seedlings (mean ± SE, $n$ = 3). Seedlings were exposed to 28 °C for three days starting on day four. The gel image depicts amplification of full-length *SR34* transcripts from cDNA of four-week-old plants of the two T-DNA insertion lines. Note that for SALK_102166 no full-length transcript of *SR34* was detected. The asterisk indicates a different T-DNA insertion line. NTC: non-template control. **(c)** Hypocotyl lengths of seven-day-old Col-0 and *afc2-1* (Salk_118114) single mutant seedlings at 17 °C and 28 °C. Seedlings were grown at 17°C for four days and then exposed to the indicated temperatures for three days ($n$ = 34-45 seedlings per genotype and temperature). **(d,e)** Temperature-dependent flowering time of Col-0, *afc1/2/3 #1* and *afc1/2/3 #2* at 17 °C, 21 °C and 28 °C measured as rosette leaf number at flowering (d) and days till flowering (e) ($n$ = 10). Plants were grown under long-day conditions at 17 °C for ten days before they were shifted to the indicated temperatures until the onset of flowering, which was defined as the day when the first flower had opened. **(f)** Representative picture of Col-0, *afc1/2/3 #1* and *afc1/2/3 #2* plants at flowering stage at 21 °C. Statistically significant differences were determined by two-way ANOVAs followed by Tukey HSD *post-hoc* tests and are indicated by different letters ($p$ < 0.05).

**Supplementary Information References**


1. Haltenhof T, *et al.* A conserved kinase-based body-temperature sensor globally controls alternative splicing and gene expression. *Molecular Cell*, (2020).

2. Labun K, Montague TG, Krause M, Torres Cleuren YN, Tjeldnes H, Valen E. CHOPCHOP v3: expanding the CRISPR web toolbox beyond genome editing. *Nucleic acids research* **47**, W171-W174 (2019).

3. Wang Z-P, *et al.* Egg cell-specific promoter-controlled CRISPR/Cas9 efficiently generates homozygous mutants for multiple target genes in Arabidopsis in a single generation. *Genome biology* **16**, 1-12 (2015).

4. Curtis MD, Grossniklaus U. A gateway cloning vector set for high-throughput functional analysis of genes in planta. *Plant physiology* **133**, 462-469 (2003).

5. Bonnot T, Gillard MB, Nagel DH. A simple protocol for informative visualization of enriched gene ontology terms. *Bio-protocol*, e3429-e3429 (2019).
